# Revisions to the *Gliophorus irrigatus* complex (Agaricales, Hygrophoraceae, *Gliophorus*, section Unguinosae) including a new waxcap, *G. alboviscidus*, from the UK but detected globally via soil eDNA, two new species from eastern North America, *G. fumosus* and *G. parafumosus*, plus a new species, *G. calunus*, and a new combination of *Hygrophorus subaromaticus* in *Gliophorus* from western North America

**DOI:** 10.1101/2025.10.22.683655

**Authors:** David J. Harries, Clare M. Blencowe, Caio A. Leal-Dutra, D. Jean Lodge, Alison H. Harrington, Ruby Bye, Zach Pearse, Stephen D. Russell, Jessica Williams, Lauren A. Ré, Gareth W. Griffith

## Abstract

Here we report the discovery of four new agaricoid fungi in the *Gliophorus irrigatus* complex of the family Hygrophoraceae. *Gliophorus alboviscidus* sp. nov. from the UK is morphologically identical to the European *G. irrigatus* (which we neotypify), except that its basidiome is white or pale Buff coloured vs brownish grey. Two new species from eastern North America, *Gliophorus fumosus* sp. nov. (provisional name *Gliophorus* sp. ‘irrigatus-IN01’), and *Gliophorus parafumosus* sp. nov (previously labelled *G. irrigatus*) resemble *G. irrigatus* s.s. in colour and morphology but their distributions are restricted to North America. Phylogenetic reconstruction revealed that these samples form distinct clades, with >10% ITS sequence divergence from European *G. irrigatus* s.s. and from each other. Though *G. alboviscidus* sp. nov. is currently known only from two locations in the UK, searches for related sequences from eDNA (environmental DNA) sequence repositories (UNITE/GlobalFungi) suggested that this species is more widely distributed in Eurasia. *G. fumosus* and *G. parafumosus* sequences from eastern North America were divergent from both European *G. irrigatus* and *G. alboviscidus*; both were more closely related to another species with a strong odour and white/Buff basidiomes from northwestern North America, *Hygrophorus subaromaticus* for which we sequenced the holotype and recombine in the genus *Gliophorus*. We also describe a new species from northwestern North America, *G. calunus* sp. nov. (provisional name *Gliophorus* sp. ‘irrigatus-CA01’), based on vouchered specimens photographed and sequenced by an paraprofessional group, CA FUNDIS. We highlight the importance of citizen scientist groups and paraprofessionals in documenting macrofungal species and their distributions via databases such as iNaturalist, Mushroom Observer and MycoMap. Further, we discuss reasons eDNA distributions are often larger than known distributions of basidiomes, including *G. alboviscidus* and *G. fumosus*.

## Introduction

*Gliophorus irrigatus* (Pers.) A.M. Ainsw. & P.M. Kirk 2013 (Agaricales, Hygrophoraceae) was first recognised by Persoon as *Agaricus irrigatus* in pine woodland (Persoon 1801), presumably near Leiden or Gottingen where he had been based up to 1801. It is likely that the specific name referred to the highly viscid pileus (“glutino adhaerens”). Fries (1821) also accepted and described *A. irrigatus* from pine woodland in “Germaniae” (reference to Persoon’s earlier discovery), as well as from grassland in “Smoland” (Småland, southern Sweden).

However, Fries also named a similar species *Agaricus unguinosus* (unguen = soft fatty or oily substance) from Scania, also in southern Sweden (Skåne), the latter considered by Persoon to be a variety of *A. irrigatus* (Persoon 1828). Whilst *A. irrigatus* clearly has priority, more recent publications have varied in their treatment of the two species. For instance, Arnolds (1990), considered the two species to be distinct, whilst Kovalenko (1988) mentions only *A. unguinosus*. However, Boertmann (1995) stated that the less viscid pileus of *Agaricus irrigatus* was simply the result of ‘desiccation and erosion’, noting also that *A. irrigatus* has priority. He later formally synonymised *Hygrocybe unguinosus* into *H. irrigata* (Boertmann 2002).

The genus *Gliophorus* was created by Herink (1959) who also recognised only *G. unguinosus* and created Sect Unguinosae to accommodate this species. However, Herink’s renaming of *G. unguinosus* was <u>invalid</u> (nom. invalid, Art. 41.4), later corrected by Kovalenko (1988). Currently *Gliophorus irrigatus* (Pers.) A.M. Ainsw. & P.M. Kirk 2013 (IF/MB#550234) is the accepted name (Kirk 2013) but the section name remains valid.

*Gliophorus irrigatus* is reported globally, with most records from Europe (Boertmann 2010) and North America (Bessette et al. 2012) but records based only on morphology also exist from Australia (Young 2005), Asia (Japan), and Africa (Kenya) (https://www.gbif.org/species/8097905). In Europe, it is most commonly found in grassland habitats but elsewhere in broadleaf and coniferous woodlands (Griffith et al. 2013; Halbwachs et al. 2018).

A pale Buff to white species originally described as *Hygrophorus unguinosus* var. *subaromaticus* (Hesler and Smith 1942) but later renamed as *Hygrophorus subaromaticus* (A.H. Sm. & Hesler) Largent was discovered in California under *Sequoia* (Largent 1985). It was distinctive in having a faint but disagreeable odour and a near-absence of clamps on hyphae (though these are also rare in *G. irrigatus*). Apart from the type specimen from 1937, there are six records from northern California (GBIF: https://doi.org/10.15468/dl.9675az; Rockefeller: https://mushroomobserver.org/307719).The stipe/pileus colour of all of these ranged from pale Buff to white and two had an unpleasant (“plastic-like” or old rubber tire) odour, one had a pleasant odour and the others had no odour. ITS sequence have been published in GenBank for four of the Californian specimens (MG926555, OR593569, PP975575, PV791511) that match the sequence we generated from the holotype (ZAP119), one published in <u>iNaturalist:264128202</u> and one in Mushroom Observer (<u>307719</u>).

Here we describe a new white/pale Buff, viscid, odourless waxcap species from the UK that is very similar in appearance (except in colour) to *G. irrigatus* but genetically quite distinct. We also describe two new species that resemble *G. irrigatus* morphologically for two of the grey-brown eastern North American clades that are phylogenetically distinct. We also show that the distributions of eDNA variants matching most of these species are wider than the distributions currently known based on sequences obtained from basidiomes, but that *G. irrigatus* s.s., which we neotypify from Denmark, is restricted to Europe. Further, for two taxa restricted to northwestern North America we recombine *Hygrophorus subaromaticus* in *Gliophorus*, and describe a new species, *G. calunus*, replacing the provisional name *Gliophorus* sp. ‘irrigatus-CA01’.

## Methods

### Morphology

Macromorphology was assessed either when freshly collected or from photographs. Capitalized colour names are from Ridgway (1912) followed by Munsell (https://munsell.com/) colour annotations by Smithe (1975), while colour names in lower case are general descriptors.

Micromorphology of European collections was assessed with a Vickers M17 microscope using a 1.30 NA objective (oil immersion) with images captured using a Canon EOS system and analysed using MICAM v1.6 (http://science4all.nl/). Micromorphology of eastern North American collections was assessed using an Olympus BH-2 microscope, an oil immersion lens (100x) and a drawing tube. A collection of *G. calunus* (<u>iNaturalist:102000323</u>) from Washington State was examined by L. Ré at 1000x with an oil immersion objective lens using a Nikon Eclipse Ci-L microscope; images captured using a Nikon Digital Sight DSFi-2. Two collections *G. calunus* from California HAY-F-012170 (holotype; <u>iNaturalist:194601188</u>) and HAY-F-000642 (<u>iNaturalist:148430507</u>) were examined by W. Cardimona at HAY using a Olympus BX53 microscope using an oil immersion objective lens.

### DNA extraction and sequencing

DNA was extracted from voucher material using a quick extraction method (Zou et al. 2017) and amplified with the primer pairs ITS8-F (5’-AGTCGTAACAAGGTTTCCGTAGGTG-3’) / ITS6-R (5’-TTCCCGCTTCACTCGCAGT-3’) (Dentinger et al. 2010) for the ITS region or LR0R (5’-ACCCGCTGAACTTAAGC-3’) / LR5 (5’-TCCTGAGGGAAACTTCG-3’) using a Bentolab thermal cycler (www.bento.bio). Cycling conditions were as follows: for ITS, 95°C for 12 min, 35 cycles [of 95°C for 20 sec, 55°C for 30 sec, 72°C for 1 min] and a final extension step at 72°C for 10 min.]; for LSU, 95°C for 5 min, 35 cycles [of 95°C for 30 sec, 50°C for 30 sec, 72°C for 1 min] and a final extension step [72°C for 10 min]. Amplification of the ITS region from the holotype of *G. subaromaticus* was achieved using a new primer (SubaromF1 (5’-GAAGGATCATTAACTGAAATTTTAGGGA-3’, based on the sequence spanning the 18S/ITS1 border of GenBank:<u>MG926555</u>) in combination with ITS4 (5’-TCCTCCGCTTATTGATATGC-3’), and using the same PCR cycling conditions as outlined for ITS amplification above. The resulting amplicons were sent for Sanger sequencing at the IBERS Translational Genomics Facility (Aberystwyth University). DNA of *Gliophorus calunus* <u>iNaturalist:102000323</u> from WA, USA was extracted using a Qiagen DNeasy Plant Pro Kit, and amplified using a BioRad Thermalcycler at U. Wisconsin-LAX. DNA sequences and chromatograms were curated and assembled using Geneious Prime (https://www.geneious.com).

US samples that were sequenced by Mycota (Plymouth, Michigan, USA; mycota.com) used a MinION workflow designed by S. Russell for use with Flongle 10.4.1 flowcells and **<u>V14 Ligation</u>** chemistry (Russell 2023). A small (rice grain-sized) piece of dried basidiome was extracted with 20 µl X-Amp DNA reagent (IBI Scientific Cat. # B47441) in strip microtubes, heated for 1 hr at 80°C in a thermocycler, followed by addition of 100 µm Molecular Biology Grade Water (IBI Scientific Cat. # IB42130). Reverse primers were uniquely tagged. Ten uniquely tagged forward (ITS1F) primers were used together with a set of 96 uniquely tagged reverse (ITS4) primers (Eurofins Genomics) designed for 96-well plates. Resulting reads were basecalled with Dorado (v0 9.1), demultiplexed with Minibar (v0.24), and consensus sequences were formed with NGSpeciesID (v0.1.2.1).The reliability and accuracy of nanopore sequencing for amplicon-based barcoding has been broadly demonstrated (e.g. (Koblmüller et al. 2024);Vasiljevic, 2021 #4485}). It relies on consensus building as an error correction mechanism, producing results that in many cases match or even exceed Sanger accuracy (Srivathsan et al. 2021). Nanopore sequencing is now routinely employed in large-scale barcoding and biodiversity initiatives worldwide (Hebert et al.).

### Assembling ITS datasets

Sequences from GenBank and UNITE databases were initially identified via BLAST searches. Additional searches were undertaken for potentially related ITS1 or ITS2 sequences derived from soil eDNA metabarcoding studies via the GlobalFungi database (https://globalfungi.com/; (Baldrian et al. 2021; Větrovský et al. 2020)) using release 5.0 (16.11.2023) of the database. Taxonomy for this version of the GlobalFungi database is based on UNITE version 10.0 (04.04.2024) (Abarenkov et al. 2024). For structured GlobalFungi searches the following method was used: The “BLAST - group results (input 1 sequence; ITS1 or ITS2)” Search type was selected, to interrogate both the “Nonsingletons (all variants)” and “Singletons (annotated variants only)” databases. The search was selected to return the top 500 BLAST (sequence variant) results. The database was search using either ITS1 or ITS2 sequences of the holo/neotypes of the six species considered in the present study. Threshold for sequence identify were set from 93% to 100% identity and geographical distributions visualised via the ‘Map” tab (Table 1).

**Table 1.** Results of BLAST searches of the GlobalFungi eDNA database using ITS1 or ITS2 sequences (at thresholds from 93% to 100% identity) from the six *Gliophorus* Sect. Unguinosae used in the present study. Numbers of soil samples in which the sequences were detected are shown in brackets. Locations of these samples are shown in the final column are shown, with location of hits >97% in bold red font.

| Species | Region | SH (1.5%) | GenBank | No. samples (No. sequence variants) at each % identity threshold |  |  |  |  |  |  | Distribution |
| --- | --- | --- | --- | --- | --- | --- | --- | --- | --- | --- | --- |
|  |  |  |  | 100% | 99.5% | 99% | 98% | 97% | 96% | 93% |  |
| *Gliophorus albobiscidus* | ITS1 | SH0910869.10FU | OP220998 | 0 | 0 | 0 | 0 | 0 | 0 | 15(39) | China, E. USA (NY,TX) |
| *Gliophorus irrigatus* | ITS1 | SH0910871.10FU | KF291084 | 7(6) | 14(165) | 15(242) | 15(272) | 18(338) | 20(350) | 24(419) | Europe, Canary Is., Canada:Newfoundland, India, USA (MN) |
| *Gliophorus fumosus* | ITS1 | SH0910862.10FU | KF291086 | 10(3) | 14(97) | 15(278) | 15(386) | 15(387) | 15(387) | 18(391) | NE America (+ MN/AZ), SE Australia |
| *Gliophorus parafumosus* | ITS1 | SH0910870.10FU | MH979262 | 4(1) | 4(2) | 4(2) | 4(2) | 4(2) | 4(2) | 4(2) | Canada:Nova Scotia |
| *Gliophorus subaromaticus* | ITS1 | SH0910861.10FU | ZAP119 | 0 | 0 | 0 | 0 | 0 | 3(4) | 5(40) | Nepal, SW China |
| *Gliophorus calunus* | ITS1 | SH0910864.10FU | PQ619233 | 0 | 0 | 0 | 0 | 0 | 0 | 11 | China, E. USA (NY,TX) |
| *Gliophorus albobiscidus* | ITS2 | SH0910869.10FU | OP220998 | 0 | 0 | 0 | 0 | 8(2)* | 183(>500) |  | China x |
| *Gliophorus irrigatus* | ITS1 | SH0910871.10FU | KF291084 | 23(16) | 27(449) | ≥27(>500) |  |  |  |  | Europe (as far east as Georgia) |
| *Gliophorus fumosus* | ITS2 | SH0910862.10FU | KF291086 | 78(21) | ≥94(>500) |  |  |  |  |  | USA (westmost OR/AZ), E. Canada,Puerto Rico |
| *Gliophorus parafumosus* | ITS2 | SH0910870.10FU | MH979262 | 5(3) | 9(57) | 65(85) | 91(>500) |  |  |  | USA (westmost CO), E. Canada, Panama |
| *Gliophorus subaromaticus* | ITS2 | SH0910861.10FU | ZAP119 | 0 | 0 | 0 | 0 | 0 | 0 | ≥10(>500) | S. China |
| *Gliophorus calunus* | ITS2 | SH0910864.10FU | PQ619233 | 0 | 0 | 0 | 0 | 0 | 0 | ≥203(>500) | China, Canada |

Additional eDNA sequences in GenBank were obtained from soil and litter collected at NEON and Long-Term Ecological Research sites and a few neighbouring parks in the USA (NCBI SRA under <u>PRJNA1327291</u>, USA National Science Foundation Grant DEB-2106130). For North America, a search of the GlobalFungi database, <u>MycoBLAST.org</u> and new eDNA data from US National Science Foundation Macrosystems Biology and NEON-Enabled Science grant (DEB-2106130), using American *G. irrigatus*-like reference sequences (MH979262 from Wisconsin and KF291086 from North Carolina) yielded 8221 eDNA sequences with >97% identity to *G. fumosus*. These were used in mapping eDNA distributions.

### Sequence alignment and phylogenetic analyses

Details of the sequences used for phylogenetic reconstruction in this study are listed in Suppdata1. The dataset containing the selected sequences was partitioned into ITS1, 5.8S and ITS2 regions. Each partition was aligned with MAFFT v7.520 (Katoh and Standley 2013), using the G–INS–i algorithm. The final alignments were curated manually with AliView v1.5 (Larsson 2014).

A maximum-likelihood (ML) phylogeny was inferred using IQ-TREE v3.0.1. ModelFinder with greedy partition merging (-m MFP+MERGE) used to select the best-fit substitution model for each partition in the final scheme. Branch lengths followed the edge-linked proportional scheme (-p), i.e., partitions shared a single set of relative branch lengths while each had its own partition-specific rate multiplier. Clade support was assessed with ultrafast bootstrap (UFBoot; 3,000 replicates with gene resampling (--sampling GENE) (Hoang et al. 2018) (Suppdata 2).

Bayesian Inference (BI) was performed using MRBAYES v3.2.7 (Ronquist et al. 2012) using two independent Markov Chain Monte Carlo (MCMC) runs (four chains each, starting from random trees). The most appropriate evolutionary models and partitioning scheme were selected from IQ-TREE’s ModelFinder results. The chains were run for 10 million generations, sampling trees every 1000 generations. After discarding the first 25% of samples as burn-in, the remaining trees were summarized into a 50% majority-rule consensus, and Bayesian posterior probabilities (BPP) were mapped to nodes. Convergence and mixing were evaluated by inspecting trace plots and effective sample sizes (ESS > 200) in Tracer v1.7 (Rambaut et al. 2018), and by MrBayes diagnostics (potential scale reduction factors approx. 1.0). Nodes were considered strongly supported when SH-aLRT ≥ 80%, BPP ≥ 0.95 and/or UFBoot ≥ 95%.

## Distribution mapping

Eurasian distributions of species in *Gliophorus* sect. Unguinosae were mapped using sequenced specimens of *Gliophorus alboviscidus* from this study and soil eDNA sequences from GlobalFungi Database matching at >97% identity and mapped using Ultimaps (map of Eurasia with countries, cropped). ITS sequence of the neotype of *G. irrigatus* was used to find eDNA sequences matching at >97% identity in the GlobalFungi Database (Větrovský et al. 2020) (Suppdata1) and mapped using Ultimaps (map of Europe with countries). Specimen-based records of preserved *G. irrigatus* having GIS coordinates were mapped using GBIF, with symbols added by hand indicating sequenced specimen records from UNITE, including GenBank records and the neotype. North American species in *Gliophorus* sect. Unguinosae were mapped based on sequenced specimens reported in iNaturalist, Mushroom Observer and Mycoportal plus matching eDNA records at >97% identity to soil sequences from US Long-Term Ecological Research sites (NCBI SRA under PRJNA1327291, USA National Science Foundation CLIMUSH grant), GenBank and soil eDNA sequence records from the GlobalFungi Database (Větrovský et al. 2020).

## Results

### Sample collection from the UK and morphological examination

In October 2021 a group of viscid, pale Buff waxcaps (tinted 1.0Y 6.0/7.0; Fig. 1A/B) was discovered on soil in unfertilised ancient grassland at Hundleton, Pembrokeshire, Wales. The morphology suggested the collection might be an albino form of a known species or an undescribed taxon. The viscid pale pileus was campanulate with Buff tints and a white margin and a white viscid stipe. The stipe was visibly marbled, a common feature of *Gliophorus* spp. The lamellae were decurrent and lacked a gelatinised edge. There was no perceptible odour. Microscopically the specimen was consistent with the description of *G. irrigatus*. Thus, the macro- and micro-morphological characters were consistent with the concept of *Gliophorus* section Unguinosae Herink.

**Fig. 1.**
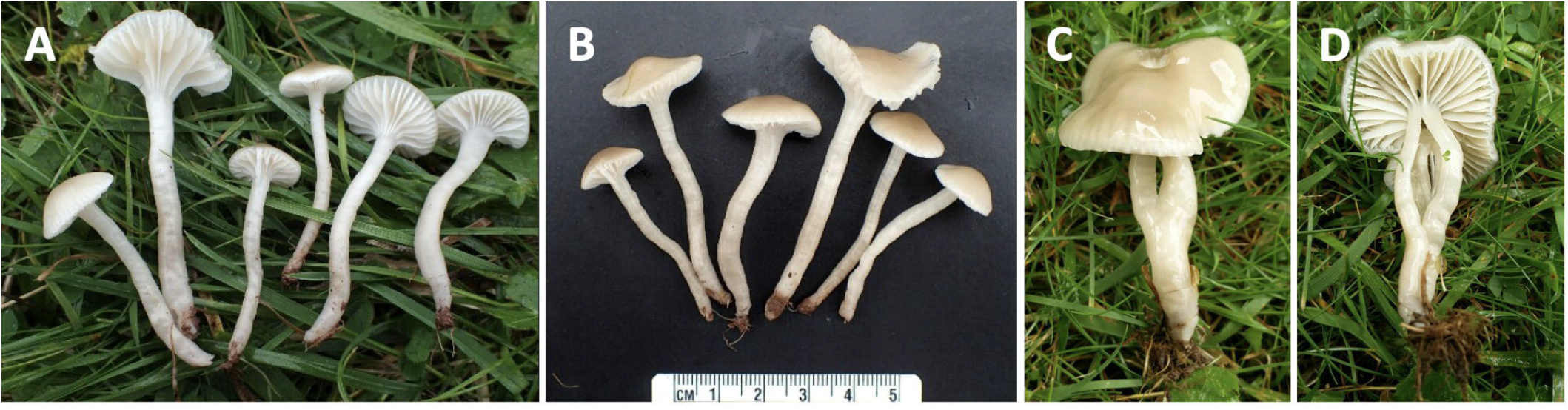
Macromorphology of *Gliophorus alboviscidus* sp. nov. basidiomes. DJH21 from Hundleton, Pembrokeshire (A,B) and CMB54 from Henfield, West Sussex (C,D).

Enquiries with members of the UK field mycology community revealed a report of a similar collection in 2019 from West Sussex (Fig. 1C,D), slightly larger and with a more strongly Buff coloured pileus/stipe, which was also microscopically consistent with the description of *G. irrigatus*. The viscid pileus and stipe and marbled appearance of the stipe clearly placed this specimen in the genus *Gliophorus*.

### Phylogenetic reconstruction

DNA sequences for the full ITS and D1/D2 domains LSU loci were obtained for both specimens. Both were identical to each other and quite distinct from other sequences of *G. irrigatus* from both Europe and North America (<91% ID across ITS region), thus excluding the possibility that they were simple albino variants.

Phylogenetic reconstruction, using sequence KY807663 (*G*. “*sciophanus*”) as the outgroup, placed these pale UK samples with high confidence in a distinct clade (Fig. 2). European specimens of *G. irrigatu*s formed a well-supported clade, quite distinct from greyish-brown eastern North American specimens, as had long been suspected (D. Jean Lodge, pers. comm. 2021). In western North America, the *Hygrophorus subaromaticus* sequences with predominantly white or pale Buff specimens formed a fourth distinct clade while a new species that included some pale Buff collections and provisionally named *Gliophorus* sp. ‘irrigatus-CA01’ formed a fifth clade. Thus, there are three unrelated clades with white to pale Buff basidiomes within the *G. irrigatus* species complex.

**Fig. 2.**
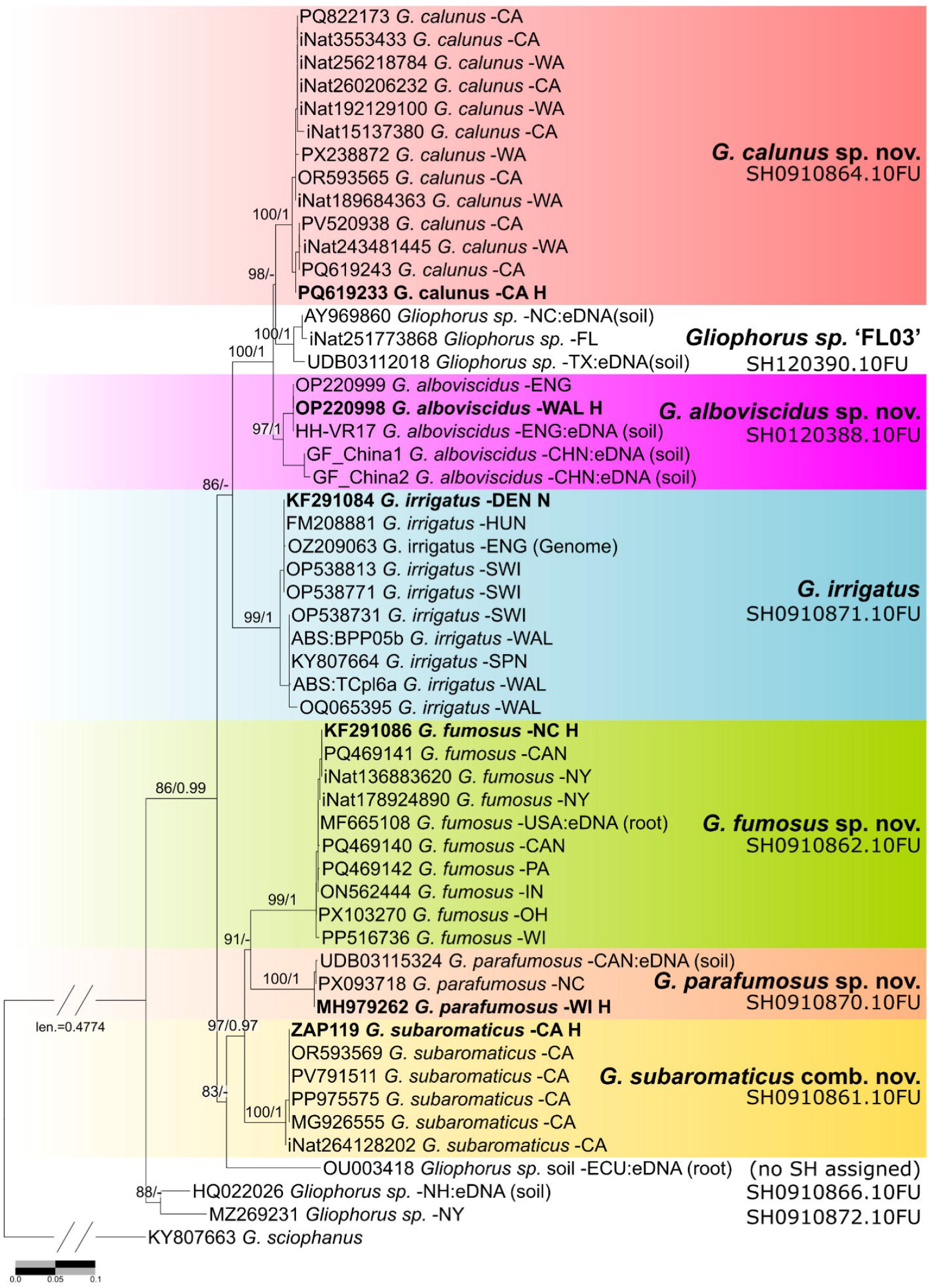
**Maximum-likelihood tree of *Gliophorus* sect. Unguinosae based on ITS** sequences, with *G. sciophanus* as the outgroup. Support values on branches are ultrafast bootstrap (UFBoot)/Bayesian posterior probabilities (BPP) and are shown only for UFBoot□≥□80 and BPP□≥□0.90. Dashes (−) indicate values below the threshold. Holotypes (H) and the proposed neotype (N) for *G. irrigatus* s.s. are indicated. UNITE Species Hypotheses (SH) at 98.5% sequence identity are shown for each clade, except for *G. alboviscidus* and *Gliophorus* sp. ‘FL03’ for which a 3% SH threshold was applied. Scale bar: nucleotide substitutions per site.

*Hygrophorus subaromaticus* resembles *G. alboviscidus* in pileus colour but belongs to a distant clade (Fig. 2). The holotype of *Hygrophorus subaromaticus* from redwood forest in northwestern US was sequenced (ZAP119) and this was 100% identical to four sequences in GenBank (MG926555, PV791511, OR593569 and PP975575; Fig. 9). Whereas the pileus colour of the type was described as buffy brown on the disc and pale Olive-buff near the white margin, two of the recently sequenced collections were entirely white (MG926555, Mushroom Observer 307719, and OR593569, <u>iNaturalist:148999043</u>), and the others were white with a Buff disc (PP975575, <u>iNaturalist:191417805</u>) or Drab Gray on the disc or overall (PV791511, <u>iNaturalist:256214946</u>. No environmental sequences close to *H. subaromaticus* clade were detected either via GlobalFungi or UNITE (Fig. 5B), suggesting that this species is restricted to the northwest coast of North America.

Seven sequenced collections from redwood forest in northern California resemble *H. subaromaticus* with pileus colours ranging from nearly white to Buff or pale greyish brown with a white margin but the former comprises a distant clade provisionally named *Gliophorus* sp. ‘irrigatus-CA01’ and described below as *Gliophorus calunus* (Fig. 6). Five additional sequenced collections were identified from Washington state under other conifer species in coastal forests. *Gliophorus calunus* can be distinguished by the blue fluorescent lamellae under UVf365 nm whereas *H. subaromaticus* lamellae remain white (refer to the key). The European clade of *G. irrigatus* s.s. is absent from North America, with most sequences derived from soils in Europe (Bahram et al. 2020; Praeg et al. 2020) (Fig. 3B).

**Fig. 3.**
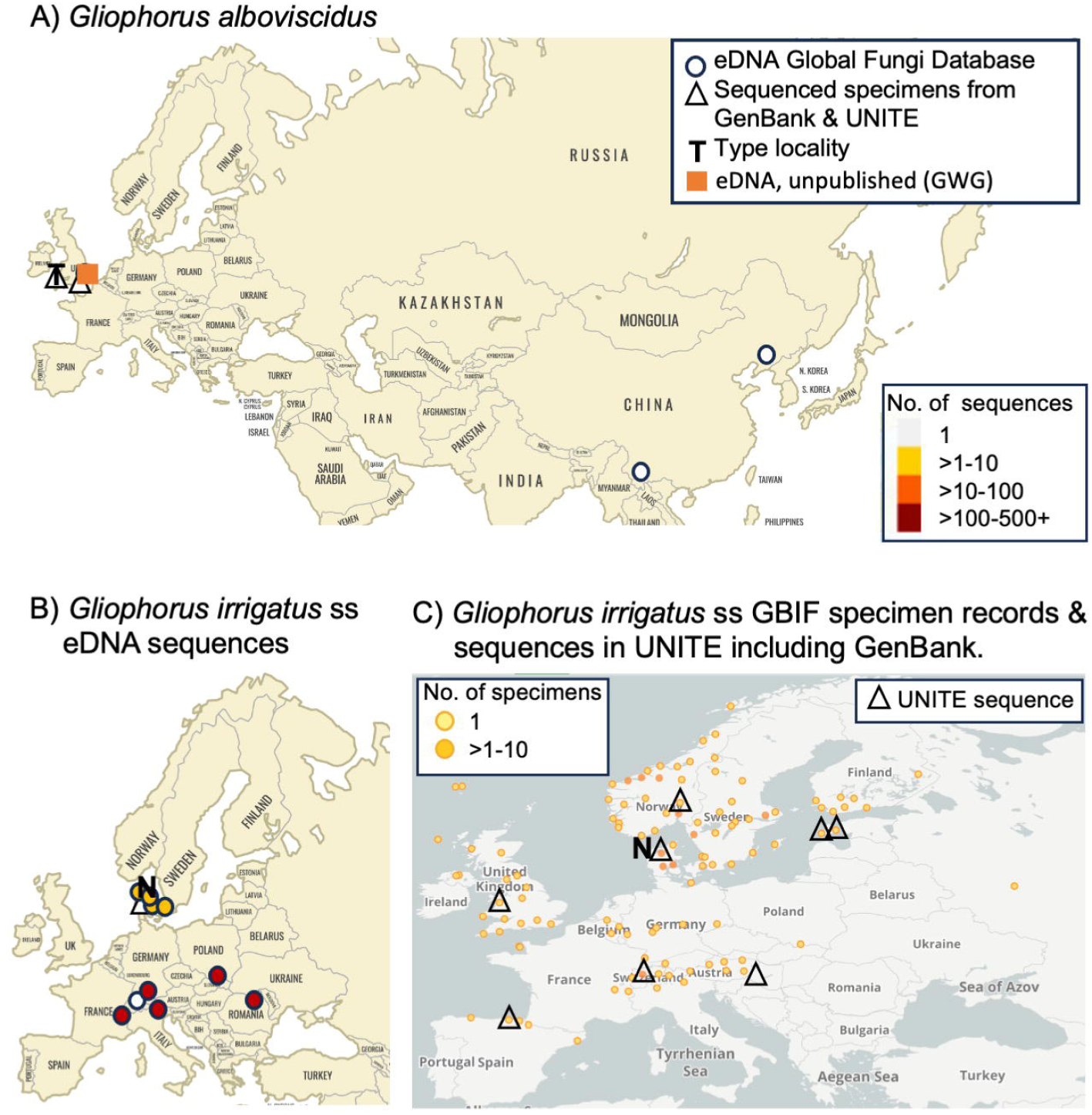
Distribution of species in *Gliophorus* sect. Unguinosae in Eurasia. A) Sequenced specimens of *Gliophorus alboviscidus* from this study and soil eDNA sequences from GlobalFungi Database matching at >97% identity. B) Previously sequenced specimen designated as the neotype of *G. irrigatus* and eDNA sequences from GlobalFungi Database (Větrovský et al., 2020) matching at >97% identity C) Specimen-based records of preserved *G. irrigatus* with GIS coordinates (<u>GBIF.org; 11 Sep 2025</u>) together with sequenced specimen records from UNITE, including GenBank records. A single occurrence of *G. irrigatus* in eastern Canada (Nova Scotia) was also detected via GlobalFungi but in not mapped on Fig. 3B.

#### Taxonomy and distributions by region

The *Gliophorus* species named or renamed in this publication all belong to Sect. Unguinosae Herink. We do not proposed to rename the Section, as this was validly published and renaming is not required under rules of nomenclature (Paul Kirk, pers. Comm. 8th Dec 2024).

### Eurasian species and distributions

Sequences falling into the *G. alboviscidus* clade (Fig. 2) were retrieved via GlobalFungi searches (ITS2 >97% identity; none with ITS1 search) from two studies in China (/Sun;(Zhang et al. 2017) (Table 1; Fig. 3A). The *G. alboviscidus* clade is geographically structured, comprised of branches with sequences from the UK and China. *G. irrigatus* s.s. was widely detected across Europe, as far east as Georgia, with ITS1 searches also detecting this species at low abundance in Newfoundland, Canada, the Faroe islands, Denmark and the Canary Islands, Spain.

### *Gliophorus alboviscidus* D.J. Harries & G.W. Griff., sp. nov

Index Fungorum: **IF559911**

#### Etymology

Refers to the white colour and viscid nature of the basidiomata.

#### Diagnosis

Basidiomata resembling *Gliophorus irrigatus* but lacking grey or brownish colours. ITS sequences strongly divergent (9.7%).

#### Holotypus

UK • Wales, Pembrokeshire, Somerton Farm near Hundleton, 51.6610, -4.9913, in undisturbed grazed grassland, 18 Oct 2021, D.J. Harries (voucher ABS: DJH21-29, [ABS=Aberystwyth University biorepository], holotype; K-M001434165, isotype) (GenBank <u>ITS:OP220998</u>; nrLSU: <u>OP221000</u>). UNITE/PlutoF <u>SH0910869.10FU</u> (1.5% threshold).

#### Description

**Pileus** 10–40 mm diam, convex to broadly campanulate, becoming applanate to subumbonate with age, margin faintly translucently striate to one third of disc, viscid, white with pale Buff tint (1.0Y 6.0/7.0) over the centre (Fig. 1A/B). **Stipe** 30–50 × 3–5 mm, cylindrical, hollow, viscid, white. **Lamellae** subdecurrent to decurrent, white. **Taste and odour** indistinct.

**Basidiospores** broadly ellipsoid or ovoid, (5.5–)6.0–8.0 × 4.5–6.0(–6.5) μm (mean 7.0 × 5.0), Q = 1.2–1.7 (mean 1.4) (type collection, 3 sporocarps, n = 70), spore print white. **Basidia** 40–50 × 5–8 μm, predominantly 4-spored. **Hymenophoral trama** sub-regular. **Pileipellis and stipitipellis** an ixotrichoderm.

#### Habitat

Recorded on soil in undisturbed grassland managed through cattle-grazing and hay cropping or of low nutrient status, regularly mown, amenity grassland, unploughed for over 50 years and not subjected to any synthetic fertiliser application.

#### Geographical distribution

Basidiomes hitherto only observed in the UK but eDNA data suggest a wider distribution across north-east Asia (Fig. 3A).

#### Other specimens examined

UK • England, West Sussex, Henfield, St. Peter’s Church, 50.9325, -0.2765, in mown grassland (cemetery) on soil, 27 Sep 2019, Clare M. Blencowe, ABS: CMB54; K-M001434166), morphologically near-identical to holotype (Fig. 1C/D) (ITS:<u>OP220999</u>).

An exact match to theITS2 sequence of this species was detected in soil eDNA from ancient grassland at Hardwick Hall, Bury St. Edmunds, Suffolk, England (52.2294,0.7023; Griffith, unpublished data; Fig. 3A).

#### Notes

By way of a common English name for this new species, we suggest Pearlescent Waxcap (*Cap Cwyr Perlaidd* in Welsh).

### Neotypification of *Gliophorus irrigatus*

In the absence of any voucher or drawing that might constitute a holotype, we propose the following:-

*Gliophorus irrigatus* (Pers.) A.M. Ainsw. & P.M. Kirk 2013 (Index Fungorum 23: 1, 2013) Basionym: *Agaricus irrigatus* Pers., Syn. meth. fung. (Göttingen) 2: 361 (1801).

Index Fungorum: **IF904305**

#### Neotypus

DENMARK • Nordjylland, Thisted, 56.92, 8.62 (600 km north of Göttingen), in dry grassland grazed by sheep, 28 Oct 2006, David Boertmann (voucher CFMR: DEN-21 [2006/70], neotype selected here ) (GenBank ITS:<u>KF291084</u>). UNITE/PlutoF: <u>SH0910871.10FU</u> (1.5% threshold).

#### Notes

It is not known whether Persoon preserved the specimen which he described as *Agaricus irrigatus*. He did not provide collection details but it is likely that the original specimen(s) was/were collected near Göttingen or Leiden. Neither Persoon’s (1801) original description of *A. irrigatus* nor Fries’ sanctioning citation (Fries 1821) for *A. irrigatus* contained any illustrations that could constitute a holotype. Enquiries about the possible existence of any relevant specimens or drawings from Christian Persoon confirmed that none were extant at Leiden (Jorinde Nuytinck, pers. comm. Oct 2024). Similar enquiries were made at Uppsala but no paintings or vouchers survive (Åsa Kruys, pers. comm.).

At the Swedish Museum of Natural History, Stockholm there is a <u>painting</u> of *A. unguinosus* by Fries (from Sunnersta ‘skog’ [forest], in Uppsala; https://herbarium.nrm.se/specimens/S0627) (Mats Wedin [NRM], pers. comm., Oct 2024). However, consistent with the opinion of the authors and others, we find no evidence that would suggest that *G. irrigatus* and *G. unguinosus* are separate species. Because *G. irrigatus* has priority, the later painting of *A. unguinosus* by Fries cannot be considered a type for *G. irrigatus*. A typical collection of *G. irrigatus* collected and photographed by J.H. Petersen in Denmark 130 km from the neotype locality is shown in Fig. 4A, and a similar collection by D. Harries in Pembrokeshire, Wales, UK (Fig. 4B).

**Fig. 4.**
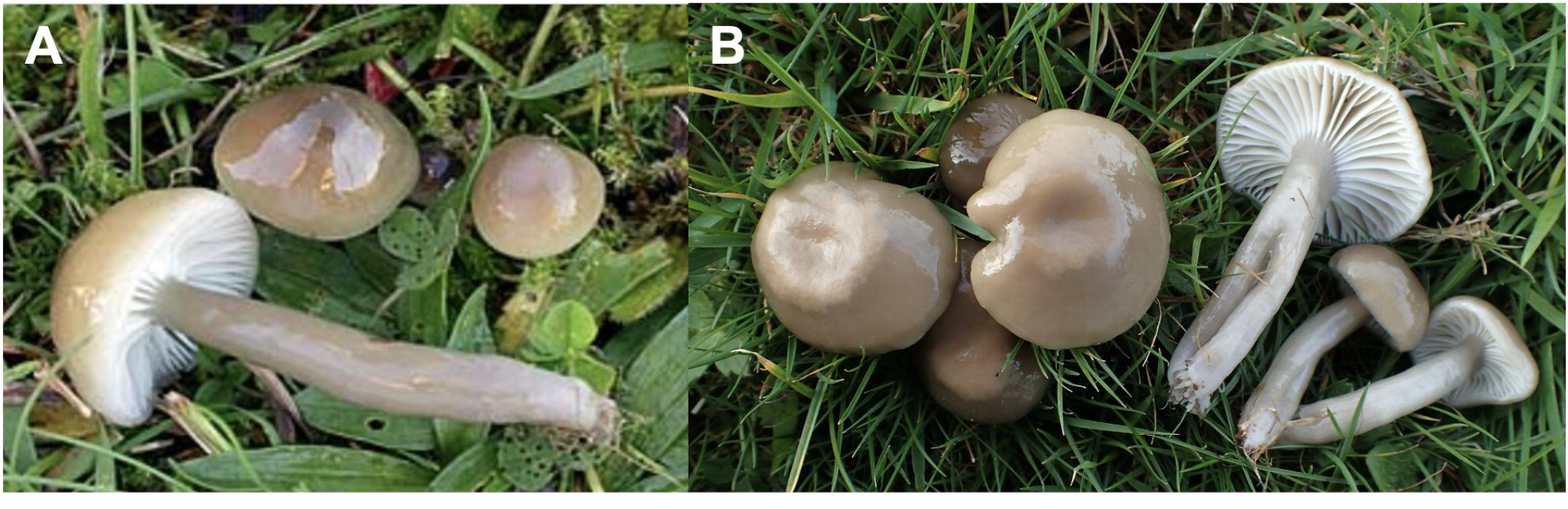
*Gliophorus irrigatus* s.s. A) Typical *G. irrigatus* basidiomes, 12 Oct 2003, Jens H. Petersen, Denmark, Høgdal, JHP-03.265, DMS-521999, 56.23676, 9.51416 (Jens H. Petersen/Mycokey) (Frøslev et al. 2025); B) *G. irrigatus* s.s. from Pembrokeshire, Wales, D. Harries (ITS: OQ065395).

**Fig. 5.**
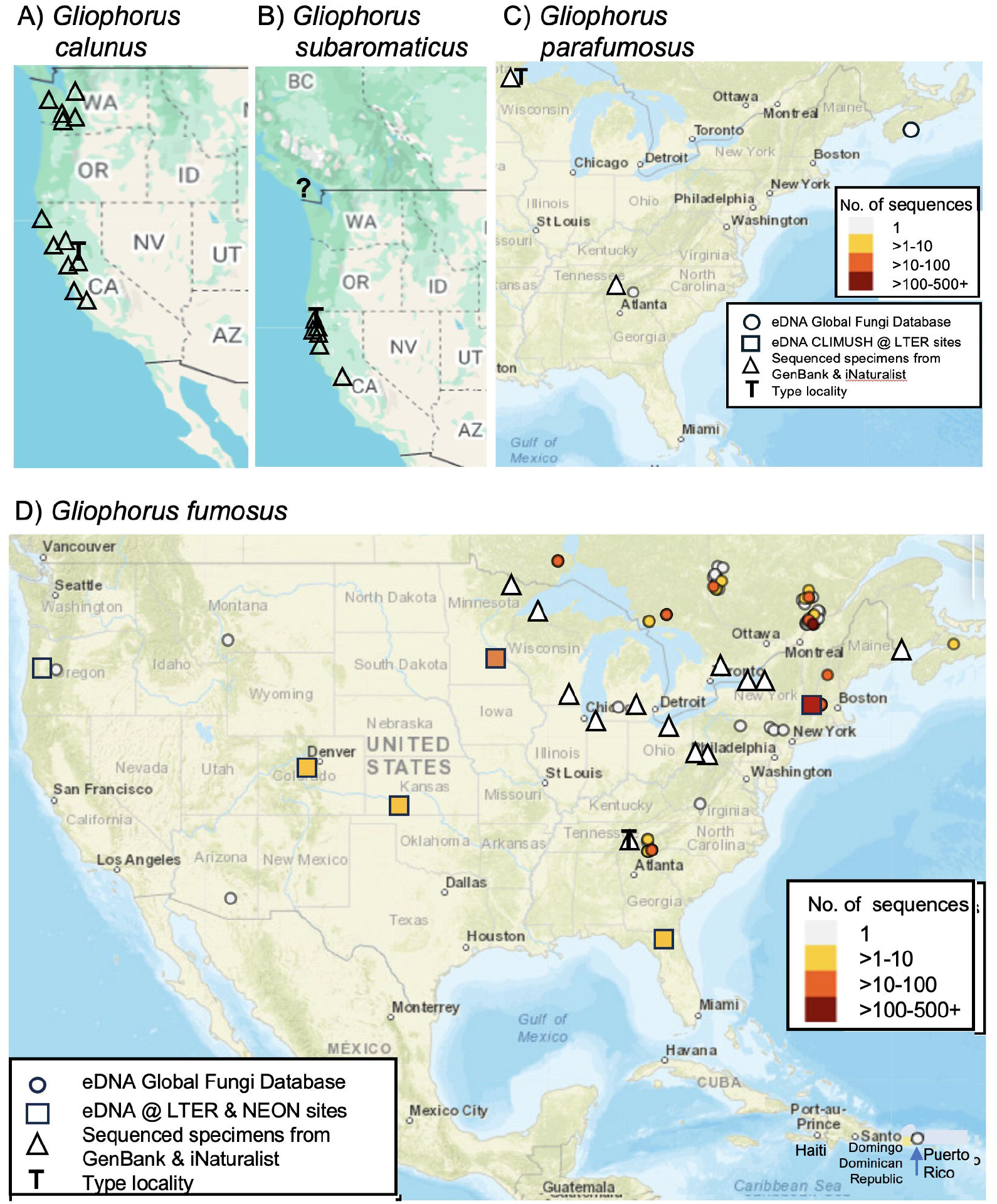
Distributions of species in *Gliophorus* sect. Unguinosae. A) Sequenced specimens of *Gliophorus calunus*. B) Sequenced specimens of *Gliophorus subaromaticus*. C) Sequenced specimens of *Gliophorus parafumosus* and two eDNA records from GlobalFungi Database (Větrovský et al. 2020). D) Sequenced specimens of *Gliophorus fumosus* and matching eDNA sequences at >97% identity to soil sequences from USA NEON-LTER sites (PRJNA1327291) and soil eDNA sequence records from the GlobalFungi Database. <u>*G. calunus* and *G. subaromaticus* were not detected in GlobalFungi database searches</u>.

The ITS sequence of this specimen was used as the representative of ‘European’ *G. irrigatus* in phylogenetic analyses of Hygrophoraceae by Lodge et al. (2014) and was already designated the <u>reference sequence</u> for this species in UNITE.

#### Distribution

Europe including the UK and southern Scandinavia according to eDNA sequences from the GlobalFungi database (Fig. 3B) and specimen records from GBIF and sequenced specimens in the UNITE database (Fig. 3C). Single occurrence of eDNA (singleton sequence) in far east of Canada.

### North American species and distributions

Sequences from several grey-brown *G. irrigatus*-like specimens, all from North America did not clearly fall in either of the clades described above. In our phylogenetic analyses, two of these collected and sequenced as part of the Great Smoky Mountains National Park All-taxa Biological Inventory (Hughes et al. 2020) were placed close to the ‘North American’ *irrigatus* clade but it is apparent that this clade comprises two main distinct clusters of sequences, one which was provisionally named *Gliophorus* sp. ‘irrigatus-IN01’, and named below as *Gliophorus fumosus* sp. nov.; it appears to be quite widespread in the eastern US (e.g GenBank: KF291086, ON562444, PP516736, PQ469140, PQ469142), and matching eDNA sequences were recovered across western parts of the USA as well (Fig. 5D). A second clade from the eastern USA and described below as *Gliophorus parafumosus* sp. nov. appears to be much less commonly found (MH979262 and PX093718; Fig. 5C). These appear strongly supported as sister clades of *G. subaromaticus* (ML 100; BPP 0.97) (Fig. 2). Additionally, a third taxon with grey-brown basidiomes was found in NY (<u>iNaturalist:30847408; MZ269231</u>), while a 91.8% similar eDNA sequence from NH remains unnamed.

A fourth North American clade with basidiomes varying from white to pale Buff, pale grey or shades of Drab (light to dark greyish brown) was reported by the California Fungal Diversity Survey (CA FUNDIS) from coastal conifer forests in northern California provisionally named *Gliophorus* sp. ‘irrigatus-CA01’, (e.g. <u>iNaturalist:194601188</u>, ITS:PQ619233; also <u>iNaturalist:148430507</u>, ITS:<u>OR593565</u>) and also found in the Puget Sound area of northern Washington State (e.g. <u>iNaturalist:189684363</u>). This species is described below as *Gliophorus calunus* sp. nov.; it has a 95.9% sequence identity to the pale Buff coloured *G. alboviscidus* from the UK despite their disjunct distributions. The fifth North American clade, *Hygrophorus subaromaticus* that we recombine in *Gliophorus* below, appears to be quite rare and hitherto recorded only from coastal redwood forests in northern California (Fig. 5B).

Searches of the GlobalFungi database (Table 1), MycoBLAST.org, and new eDNA data from US National Science Foundation Macrosystems Biology and NEON-Enabled Science grant (DEB-2106130)(Caiafa et al. 2025), with American *G. irrigatus*-like reference sequences (MH979262 from Wisconsin and KF291086 from North Carolina) yielded a larger number (8221) of matching sequences mostly linked to five studies from the USA and Canada with additional eastern North American records from iNaturalist, suggesting that this clade is restricted to North America (Fig. 5C-D).

### *Gliophorus calunus* L.A. Ré, D. Tighe, W. Cardimona, M. Hackney & Leal-Dutra, sp. nov

Index Fungorum: **IF904063**

#### Etymology

Cal – for California where the holotype and most of the additional collections were found by the California Fungal Diversity Survey, and unus-Latin for one, referring to the number 01 in the provisional name for this species.

#### Diagnosis

Pileus glutinous, margin usually inrolled, Pearl Gray, Light Drab to Drab, Dark Drab, Cinnamon or Tawny Olive at center, usually paler toward margin. Stipe viscid with horizontal bands of contrasting translucence. Resembling *Gliophorus subaromaticus* from the same northwestern North American coast to coastal range conifer forests but differs in pileus center usually with Drab tones vs Buff to Olive-buff or buffy brown tones, lamellae bright blue vs white under UVf365 nm, pileus margin usually inrolled vs straight, and lacking an odour vs usually with a disagreeable odour.

#### Holotypus

USA • California, Mendocino Co., Jackson Demonstration State Forest, 39.4045, -123.7537 (accuracy 12 m), ca. 140 m asl, 22 Dec 2023, D. Tighe, (voucher HAY:F-012170 [California State University East Bay Fungarium], <u>iNaturalist:194601188</u>, holotype) (GenBank ITS:<u>PQ619233</u>). UNITE/PlutoF: <u>SH0910864.10FU</u> (1.5% threshold).

#### Description

**Pileus** 6-24 mm diam, convex to broadly convex with an incurved margin, becoming plane or nearly so, sometimes with an umbo, colour Pearl Gray in type, Light Drab to Drab, Cinnamon or Tawny Olive, usually paler toward margin, strongly viscid to glutinous especially when young, margin often translucent-striate. **Stipe** 20–60 × 3-9 mm, often concolorous with the pileus, slimy viscid, horizontal bands of contrasting translucence (marbled), glabrous, equal or slightly tapered at base, some slightly curved or nodulose; context hollow. **Lamellae** sinuate or slightly arcuate, white, sometimes tinted pale grey or drab, 1–2(–3) lengths lamellulae inserted, bright blue under UVf365 nm. **Taste and odour** indistinct.

**Basidiospores** hyaline, ellipsoid, (6.7–)7.5–9.6(–10.8) × (4.1–)4.4–5.5(–6.5) µm, mean Q = 1.7–1.8 in the holotype and <u>iNaturalist:148430507</u> from northern California, (6.1–)6.5–8.6(– 9.6) × (4.1–)4.3–5.2(–5.6) µm, mean Q = 1.6 in <u>iNaturalist:1102000323</u> from Olympia in northern Washington. Basidia 4-sterigmate, with typical (non-toruloid) basal clamp connections. **Pileipellis and stipitipellis** an ixotrichoderm, with slender embedded hyphae lacking toruloid clamp connections.

#### Habitat

Holotype growing on a mossy sandbank among grasses, sedges and cattails, near *Vaccinium ovatum* and *Notholithocarpus densiflorus*. Typically found in coastal Cupressaceae-dominated forests (coastal redwood, *Sequoia sempervirens*, and western red cedar, *Thuja plicata*), often mixed with hardwoods and other conifers (western hemlock and Douglas fir).

#### Geographical distribution

Occurs in coastal forests of western North America, recorded from northern California and western Washington state, USA but likely occurring throughout coastal Cupressaceae-dominated forests in western North America, especially *Thuja plicata*. (Fig. 5A).

#### Additional specimens examined

USA • (1) California, Humboldt Co., Eureka Area, Headwaters Forest Reserve, Headwaters Forest Trail, 40.6897, -124.1309, ca. 45 m asl, under redwood (*Sequoia*) and alder (*Alnus*), Feb. 2023, M. Hackney, iNaturalist:148430507, ITS:<u>OR593565</u> (HAY-F-000642); (2) Washington, Thurston, County, N of Olympia, W of South Bay, 47.0738, -122.9012, ca. 10– 20 m asl., 22 Nov 2021, L. Ré & S. Hickey (<u>iNaturalist:102000323</u>).

#### Additional sequenced specimens with photographs

USA • (1) California, Humboldt Co., Trinidad, Strawberry Rock Trail, 41.0790, -124.1313 (187 m accuracy), ca. 190 m asl, 17 Nov 2023, N. Siegel CSALVA 37, northern Franciscan (coastal) redwood forest, (NS1093, UCSC); (2) Marin Co., Bolinas, Bolinas Fairfax Rd., 37.9404, -122.6577, ca. 45 m asl, under coastal redwood (*Sequoia sempervirens*), Tanoak (*Notholithocarpus densiflorus*), and *Vaccinium* spp., 6 Jan 2024, D. Tighe, <u>iNaturalist:195986180</u>, ITS:PQ619243, (HAY-F-012180); (3) ibid., Mendocino Co., Jackson Demonstration State Forest, near Caspar Orchard Rd., 39.3606, -123.7787, ca. 110 m asl, under coastal redwood (*Sequoia sempervirens*), western hemlock (*Tsuga heterophylla*), evergreen huckleberry (*Vaccinum ovatum*), and tanoak (*N. densiflorus*),19 Jan 2025, D. Lyons, <u>iNaturalist:259023907</u>, ITS:PV520938 (HAY-F-013708); (4) ibid, San Mateo Co., SW of Redwood City, 37.4700, -122.27 (28.4 km accuracy), Jan 2025, Y.-M. Wang, <u>iNaturalist:260206232</u>; (5) ibid., Santa Cruz Co., Santa Cruz, Henry Colwell Redwoods State Park, 37.0195, -122.0436, ca. 160 m asl, 13 Feb 2023, D. Lyons, <u>iNaturalist:148743563</u>, ITS:PQ822173 (OMDL11). (6) Washington, King Co., Auburn, 188th Ave. SE, 47.2675, - 122.0914, ca. 45 m asl, 31 Oct 2023, Y.-M. Wang, <u>iNaturalist:189684363</u> (PSMS1465); (7) ibid., Mason Co., near Little Skookum Inlet and SE Lynch Rd., 47.1480, -123.0630, ca. 10– 20 m asl, 2 Dec 2024, S. Ness, <u>iNaturalist:253949042</u> (OLY-0918); (8) ibid., Thurston, County, N of Olympia, W of South Bay, 47.06976, -122.89716, ca. 20 m asl, 25 Nov 2023, B. Funkhouser, <u>iNaturalist:192129100</u>; (9) ibid., Boston Harbor, near Boston Harbor Rd. NE and Bromley Ln NE, 47.13150, -122.89779, 8 m accuracy, ca. 10 m asl, 25 Dec 2024, M. Koons, <u>iNaturalist:256218784</u>.

#### Notes

*Gliophorus calunus* resembles *G. subaromaticus* but differs in pileus centre usually with Drab tones vs Buff to Olive-buff or buffy brown tones, lamellae bright blue vs white under UVf365 nm, pileus margin usually inrolled vs straight, and lacking an odour vs usually with a disagreeable odour. Both species occur in northwestern North American coastal forests dominated by Cupressaceae, with *G. calunus* growing from soil beneath both *Sequoia sempervirens* and *Thuja plicata* whereas *G. subaromaticus* has hitherto only been recorded under *S. sempervirens. G. calunus* appears to be a more widespread and abundant species (Figs. 5A,B). The pileus of *G. fumosus*, hitherto only collected in eastern North America but with eDNA sequences extending into western North America (Fig. 5D), is usually tinted more greyish brown and differs in having a straight margin. Coastal conifer forests where *G. calunus* likely occurs in northwestern North America are currently threatened by heavy logging. We suggest California Slimy Waxcap as the common English name for this new species.

### *Gliophorus fumosus* D.J. Lodge, S.D. Russell & Leal-Dutra sp. nov

Index Fungorum: **IF902428**

**Fig. 6.**
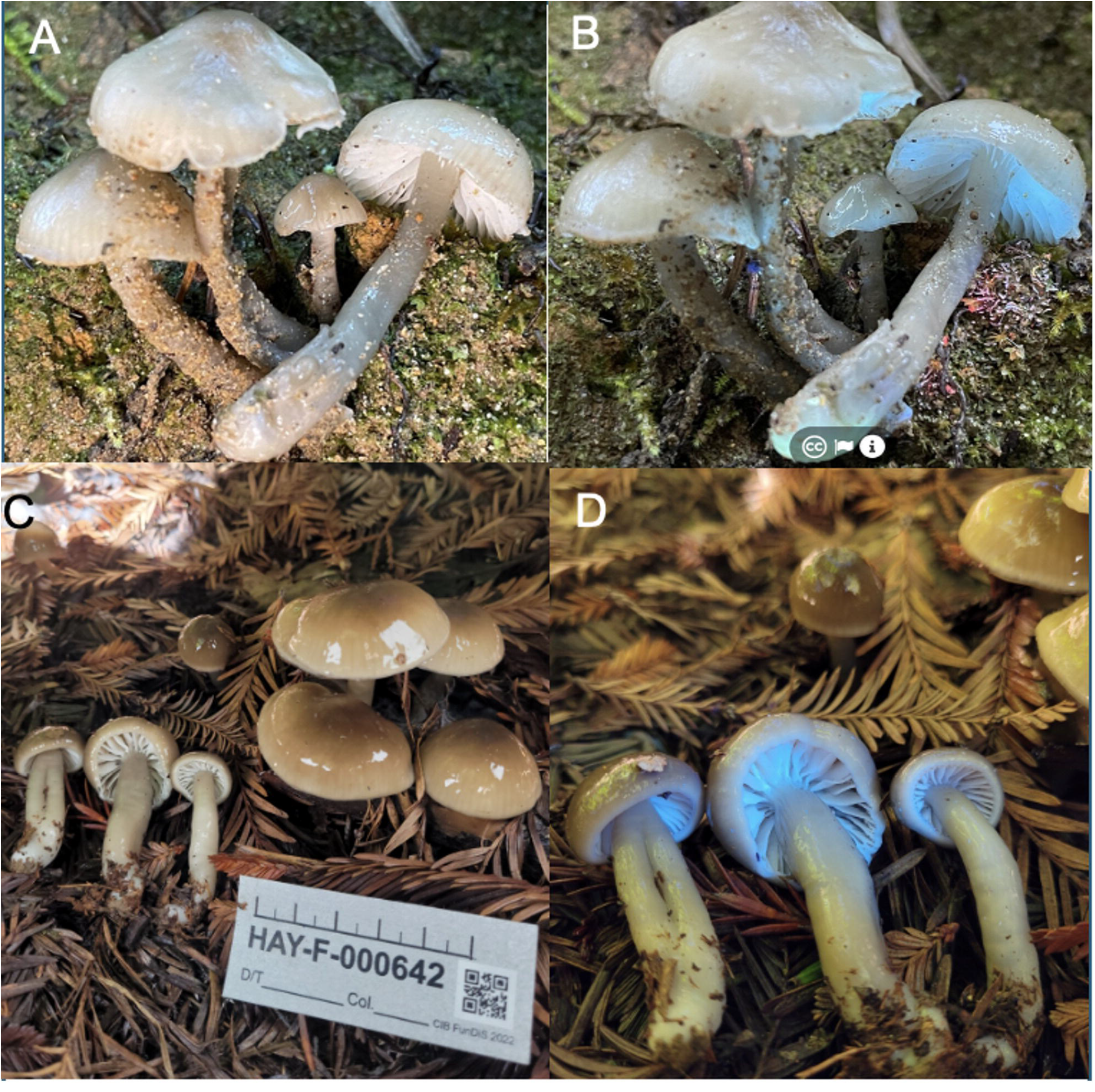
Macromorphology of *Gliophorus calunus* from northwest USA. A-B) Holotype HAY-F-012170 <u>iNaturalist:194601188</u> (ITS:PQ619233), Damon Tighe; A) in sunlight; B) in UVf365 nm light. California, Medacino Co., near Fort Bragg, 21 Dec 2023. C-D) HAY-F-00642 <u>iNaturalist:148430507</u>), California, Humboldt Co., near Eureka. Mandy Hackney, Feb 2023; C) in sunlight; D) in UVf365 nm light.

**Fig. 7.**
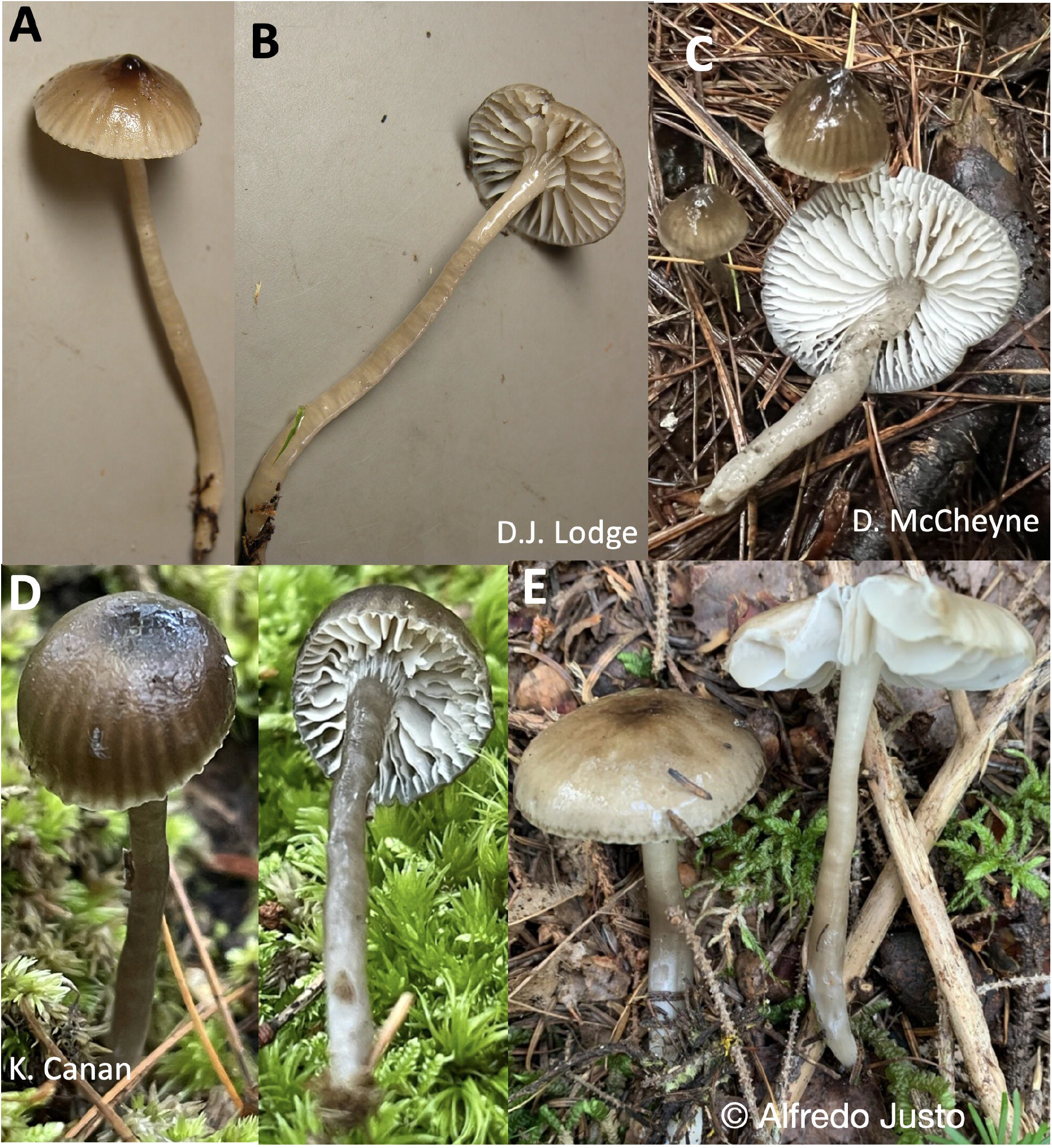
Macromorphology of *Gliophorus fumosus* basidiomes. A,B) Holotype of *G. fumosus*, DJL05NC50 from USA: North Carolina, Haywood County, Great Smoky Mountain National Park, Cataloochie, 12 Aug 2005, E.B. Lickey & D.J. Lodge (ITS:KF291086); C) USA, New York, Rome, 27 Sept 2022, D. McCheyne, <u>iNaturalist:136883620</u> (ITS:DMc055); D) USA, Wisconsin, La Pointe, Big Bay State Park, 19 Sep 2023, Kyle Canan <u>iNaturalist:183983484</u>, (ITS:PP516736); E) Canada: New Brunswick, Queens Co., Canning, 8 Sep 2022, A. Justo, <u>iNaturalist:134364174 (ITS:PQ469140</u>).

#### Etymology

Fumosus = smoke, referring to the type locality in the Great Smoky Mountains National Park and to the smokey colour of the stipe.

#### Diagnosis

Basidiomata resembling *Gliophorus irrigatus* but basidiospores shorter (5.6–7.4 vs 6.5–9) and hitherto recorded only in North America.

#### Holotypus

USA • North Carolina, Haywood Co., Great Smoky Mt. National Park, Cataloochie Cove, near Rough Fork & Big Fork Ridge trails, 35.6164 -83.1208, in forest, 12 Aug 2005, E.B. Lickey & D.J. Lodge (voucher TENN-F-061868 [DJL05NC50], holotype) (GenBank ITS:<u>KF291086</u>). UNITE/PlutoF:<u>SH0910862.10FU</u> (1.5% threshold).

#### Description

**Pileus** 10–35 mm diam, shape variable among and within collections, narrowly to broadly campanulate with a mammalate umbo when young in the type and <u>iNaturalist:145165934</u> and <u>iNaturalist:178924890</u>, often parabolic-umbonate or convex-hemispheric with a low umbo, becoming broadly parabolic or broadly convex with age with or without a low umbo, margin straight, translucently striate one third to two thirds to centre, glutinous or viscid, Drab (9.0 YR 5.5/2.5) or greyish brown with darker striae and Dark Grayish Brown (6.0 R 2.0/1.0) centre (Fig. 7). **Stipe** 30–60 × 2–5 mm, cylindric, hollow, often tapered at base and flared at apex, rarely caespitose; surface Smoke Gray (2.5 Y 6.0/2.4 to 5.0 Y 7.0/2.0), drab grey or Drab (9.0 YR 5.5/2.5), glutinous to viscid, hyphae with small clamps. **Lamellae** 5–10 mm broad, sinuate with a small to large decurrent tooth, white, pale grey toward pileus, regular, 1–2 lengths of lamellulae inserted, edges concolorous, slightly wavy. Lamellae of <u>iNaturalist:290749159</u> emitted an intense blue light under UVf365 nm. **Taste and odour** indistinct. **Basidiospores** broadly ellipsoid or ellipsoid, 5.6–7.2(–7.4) × 4–5.6 µm (mean 6.7 × 4.6), Q = 1.3–1.6(–1.7), (mean 1.5) (type collection n = 20). **Basidia** 35–82 × 7.2–15.2 µm, 4-spored with medallion clamps at base. **Hymenophoral trama** subregular, comprised of cells 19.2–82 × 8.8–15.2 µm, gelatinization absent, conducting elements abundant in context at the lamellar edge. **Pileipellis and stipitipellis** an ixotrichoderm.

#### Habitat

Recorded on forest soil, with one eDNA record detected in Fagaceae roots (Arkansas, USA). <u>iNaturalist:290749159</u> was collected between a bog and pond under *Quercus* sp. and *Nyssa sylvatica*, with abundant Vaccinieae in the understorey. Habitats covered in eDNA studies include prairie and alpine habitats extending to western USA in addition to eastern forests.

#### Geographical distribution

Basidiomes known only from eastern North America but eDNA data suggests distribution as far west as Arizona, Colorado and Oregon (Fig. 5D).

#### Other specimens examined

USA • (1) Indiana, Porter Co., North Mineral Springs, Indiana Dunes National Lakeshore, Dune Acres, Cowles Bog Trail, 41.6495, -87.0744, 4 Jul 2017, S.D. Russell, MM6069 (PUL 00035147; F 20317), (ITS:<u>ON562444</u>); (2) Ohio, Summit Co., Green, Nimisila Reservoir Metro Park, 40.9369, -81.5155, 18 Jun 2025, Jessica Williams, 2025SMP20, <u>iNaturalist:290749159</u> (ITS:<u>PX103270</u>), KEF015 as “*Gliophorus ‘irrigatus*-IN01’”.

#### Additional Records Based on Photographs and ITS Sequences

Sequenced photographic records posted on iNaturalist are all <1% divergent from the holotype sequence (all within <u>SH0910862.10FU</u>), provisionally named ‘irrigatus-IN01’. CANADA • New Brunswick, Queens, Jemseg Grand Lake Watershed (45.9458, -66.1106, 8 Sep 2022, A. Justo [voucher at NBM]), <u>iNaturalist:134364174</u>, (ITS:<u>PQ469140</u>). USA • (1) Wisconsin, Ashland Co., Apostle Islands LTA, Madeline Island, 46.7922, -90.6656, 19 Sep 2023, K. Canan [voucher OMDL01567]), <u>iNaturalist:183983484</u>, (ITS:<u>PP516736</u>); (2) Pennsylvania, Huntingdon Co., Alan Seeger Natural Area, 40.6944, -77.7552, 18 Aug 2023, J. Plischke, voucher JP23-0492, <u>iNaturalist:193184758</u> (ITS:<u>PQ469141</u>); (3) Pennsylvania, Somerset Co., Sequanota Lutheran Conference Center and Camp Estate (40.17824, - 79.10461, 2 Sep 2023, J. Plischke [voucher N23-1072]), <u>iNaturalist:181314311</u> (ITS:<u>PQ469142</u>); (4) New York, Oswego Co., Pulaski, 43.5510, -76.2047, 17 Aug 2023, P. DeSanto, voucher CM23-80169, <u>iNaturalist:178924890</u> (ITS:<u>CM23-80169</u>); (5) New York, Oneida Co., Rome, Sand Plains Unique Area (43.23238, -75.56605, 27 Sep 2022, D. McCheyne, voucher DMc055, <u>iNaturalist:136883620</u>. (ITS:<u>DMc055</u>).

#### Additional Environmental Sequences

USA: Arkansas, clone OTU_368, from (ectomycorrhizal) Fagaceae roots, (ITS2:<u>MF665108</u>). DNA of this species was detected in a range of woodland, prairie and alpine habitats in North America (Fig. 5D; 617sequences deposited at NCBI SRA under <u>PRJNA1327291</u>).

#### Notes

The macroscopic characters are not diagnostic for this species. Macroscopic characters such as the presence of an umbo varies among collections and among specimens within a collection, and the microscopic characters do not vary distinctly among species in this complex. The basidiospores are slightly shorter than those of *G. irrigatus* s.s. from Europe as noted in the diagnosis. Microphotographs of basidia and basidiospores of J. Williams 2025SMP20 are posted on <u>iNaturalist:290749159</u> (ITS:<u>PX103270</u>). ITS sequences, however, are diagnostic and differ by about 10% between clades. The bright blue appearance of lamellae under UVf365 nm illumination is distinctive but appearance of other eastern species under UV light is lacking. We suggest Smoky Waxcap as the common English name for this new species.

### *Gliophorus parafumosus* D.J. Lodge, S.D. Russell & G.W. Griff. sp. nov

Index Fungorum: **IF904176**

**Fig. 8.**
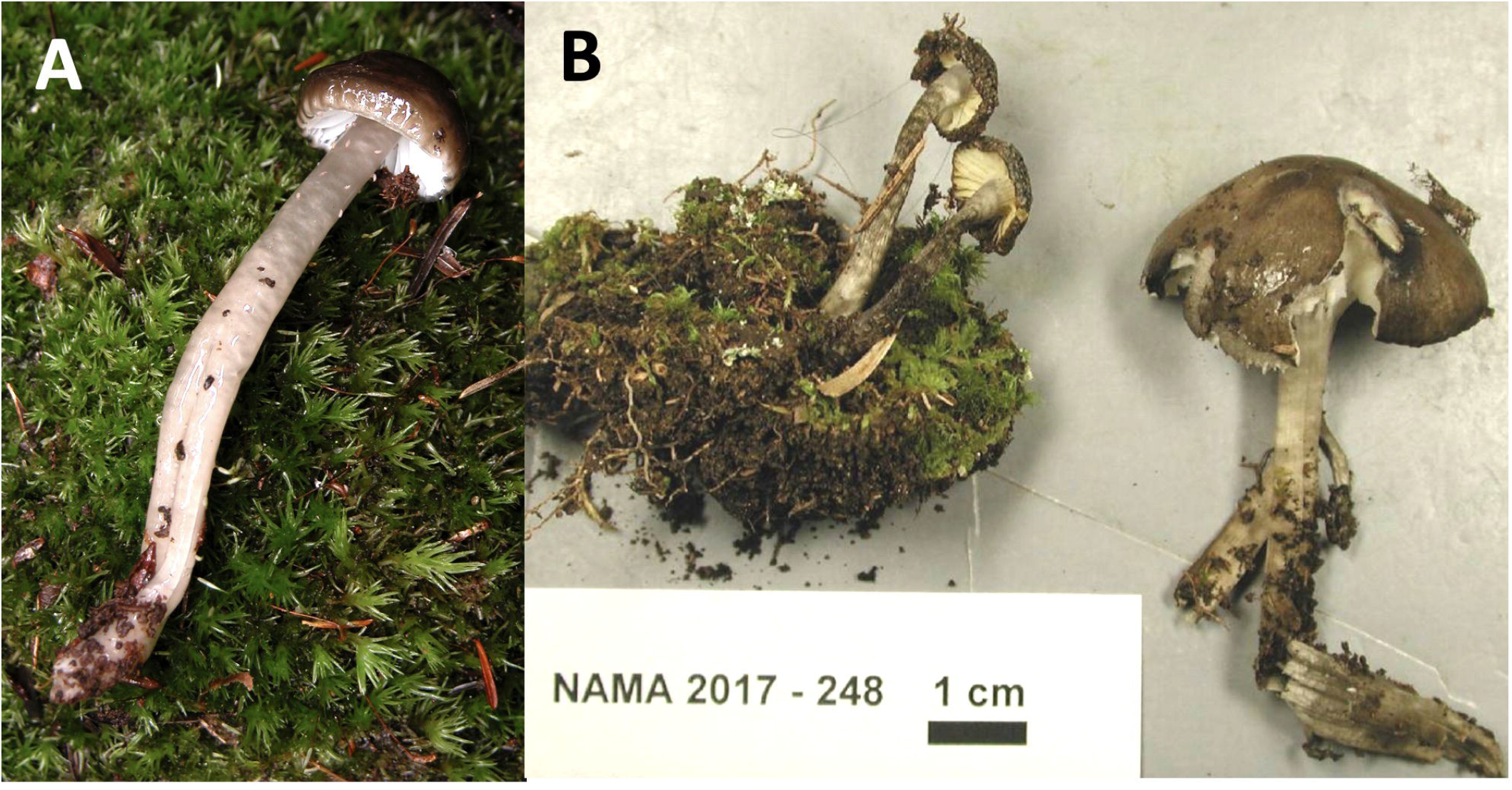
Macromorphology of *Gliophorus parafumosus* basidiomes. A) *G. parafumosus* holotype NC-91 (ITS:PX093718). USA: North Carolina, Swaine County, Great Smoky Mountain National Park, Mingas Mill, near Graveyard, (Lat/long: 35.5219, -83.3096; 360 m asl), 15 Aug 2005, D.J. Lodge. B) *G. parafumosus* NAMA 2017-248 (ITS:MH979262). USA: Wisconsin, Bayfield, Big Bay State Park (Lat/long: 46.7976,-90.6697), 8 Sep 2017, S. Wunderle, Mycoportal:<u>C0348314F</u>.

#### Etymology

Parafumosus; para= close by and fumosus = smoke, referring to its phylogenetic proximity, as well as its similarity in both morphology and distribution to *G. fumosus*.

#### Holotypus

USA • Wisconsin, Bayfield, Big Bay State Park, 46.7976, -90.6697, 8 Sep 2017, S. Wunderle, (voucher F:<u>C0348314F</u> [NAMA 2017-248] [F=Field Museum of Natural History biorepository]) (GenBank ITS:<u>MH979262</u>). UNITE/PlutoF: <u>SH0910870.10FU</u> (1.5% threshold).

#### Diagnosis

Basidiomata similar macro- and microscopically to *Gliophorus fumosus* and having the same geographic range but differing in ITS sequence and by having a darker pileus umbo (Burnt Umber to Fuscous vs Warm Sepia to Dark Drab) and an inrolled rather than straight pileus margin.

#### Description

**Pileus** 12–35 mm diam, parabolic when young, becoming broadly parabolic with age, margin inrolled and slightly sulcate when young and less so with age, surface strongly viscid, colour Burnt Umber in centre when young, Fuscous with age, Drab on margin when young and overall with age. **Stipe** 25–75 × 3–8 mm, broader in lower 1/3, some compressed, hollow, surface strongly viscid, Smoke Gray (2.5 Y 6.0/2.4 to 5.0 Y 7.0/2.0), or tinted drab grey. **Lamellae** White or pale Cream Color, sinuate or adnate with a decurrent tooth, 4–8 mm broad, margin wavey, one length of lamellulae inserted. **Taste and odour** indistinct. **Basidiospores** ovoid, hyaline, (5.2–)5.6–4(–8) × 4×5.2 µm. mean 6.4 +/- 0.76 × 4.6 +/- 0.39 µm, with or without guttulate contents, inamyloid. **Basidia** 4-sterigmate, 28–37 × 4.8–8 µm, with a small basal clamp. **Subhymenium** 10–12 µm deep, of interwoven hyphae 2.4–4.8 µm wide. Lamellar context subregular, comprised of swollen elements 30–80 × 8.8–18.5 µm rarely with small clamp connections; lamellar edge not gelatinized. **Cheilocystidia** absent. **Pileus context** of swollen hyphae 80–120 × 8–20 µm, with pale brown contents in KOH, and small, rare clamp connections. **Pileipellis** an ixocutis, hyphae embedded in gelatinous matrix 1.2–2.4 µm, rarely with small clamps. Stipe context hyphae parallel, 5.6–16 µm diam with pale brown contents, slightly constricted at septa, lacking clamp connections. **Stipitipellis and ixotrichoderm**, hyphae embedded in gelatinous matrix 1.2–3 µm diam with few small clamps.

#### Habitat and Distribution

Hitherto recorded only in North America. Growing in moss in moist, mixed deciduous forests of eastern North America (Fig. 5C).

#### Other specimens examined

USA • (1) North Carolina, Swaine County, Great Smoky Mountain National Park, Mingas Mill, near Graveyard, (35.5219404, -83.3096265) 360 m asl, 15 Aug. 2005, D.J. Lodge, NC-91, (ITS:PX093718)

#### Additional Environmental Sequence

Canada • Halifax, UNITE soil sample <u>TUE002679, UDB03115324</u>, 8 Nov 2019 (Tedersoo et al. 2014),

#### Notes

Differs from *G. fumosus* in having a pileus margin that is distinctly inrolled and sulcate-striate, in contrast to *G. fumosus* (and *G. irrigatus*) has/have a straight or rarely slightly in-rolled gluten on the margin and is either not striate or only translucent-striate - not sulcate. Based on numbers of sequence-confirmed vouchers, *G. parafumosus* appears to be less common that *G. fumosus*. The microscopic observations were from the type collection deposited at F. The collection NC-91 was not annotated because it was immature, and the remaining piece was not deposited at TENN after removing part for sequencing. We suggest False Smoky Waxcap as the common English name for this new species.

### *Gliophorus subaromaticus* (A.H. Sm. & Hesler) Rockefeller, G.W. Griff. & D.J. Harries, comb. nov

Index Fungorum**: IF901873**

**Fig. 9.**
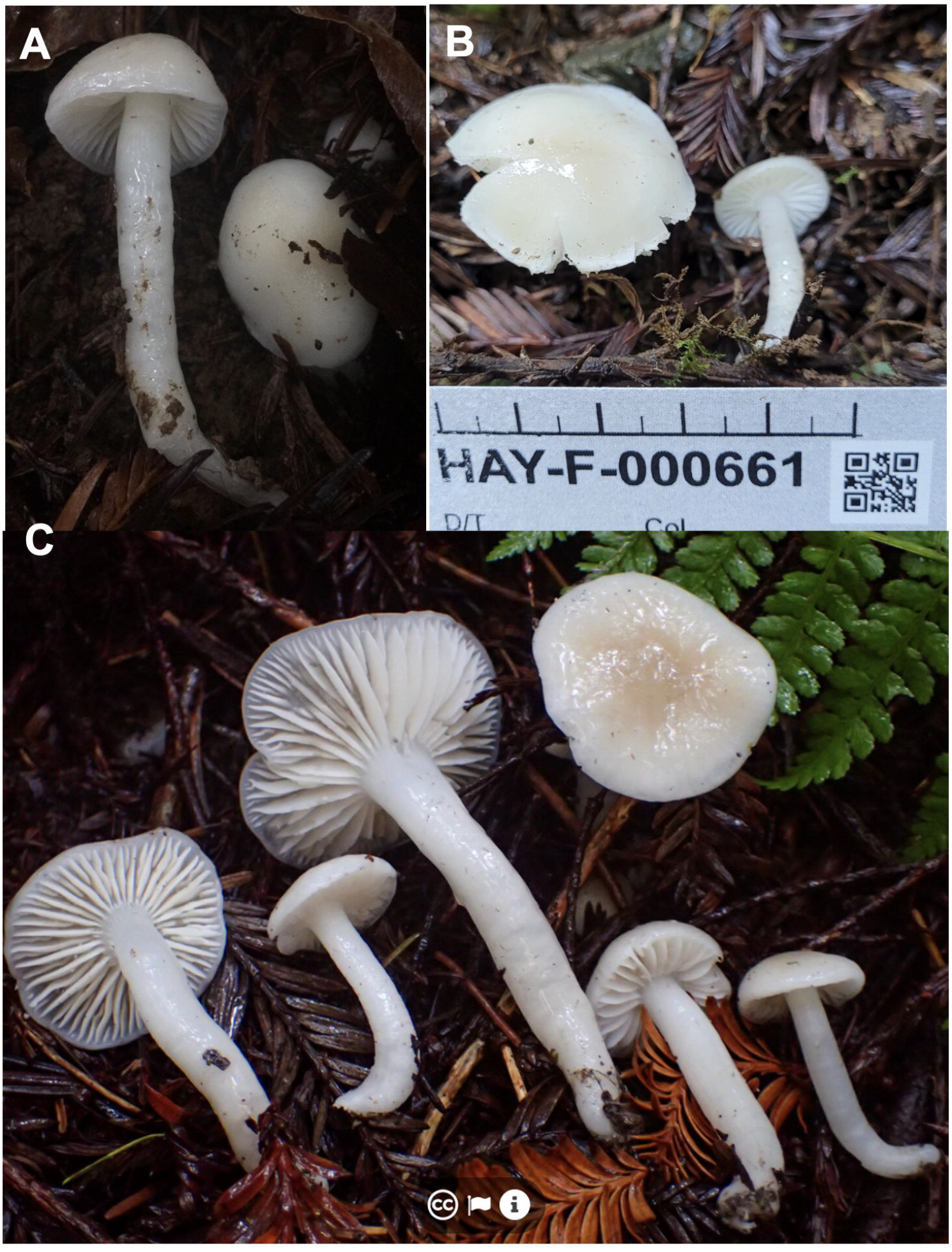
Macromorphology of *Gliophorus subaromaticus* basidiomes. A) *G. subaromaticus* **USA**, California: (1) Mendocino Co., Jackson State Forest”, (39.3890,-123.68460, 26 Jan 2018, A. Rockefeller [voucher AR-2018a]), <u>iNaturalist: 9640562</u> (ITS: <u>MG926555</u>); B) Humboldt Co., Eureka Area, Arcata Community Forest, near *Sequoia sempervirens* (40.88326, -124.06329, 16 Feb 2023, Mandy Hackney [voucher HAY-F-000661]), <u>iNaturalist: 148999043</u> (GenBank ITS sequence: OR593569);C) Humboldt Co., Eureka Area, McKay Community Forest, near *Sequoia sempervirens* (40.7501, -124.1446, 18 Nov 2023, Dean Lyons [voucher HAY-F-002303]), <u>iNaturalist #191417805</u> (ITS :<u>PP975575</u>). UNITE/PlutoF: SH0910861.10FU (1.5% threshold).

*Gliophorus subaromaticus* (A.H. Sm. & Hesler) Rockefeller, G.W. Griff. & D.J. Harries, comb. nov.

Basionym: *Hygrophorus unguinosus* var. *subaromaticus* A.H. Sm. & Hesler, *Lloydia* 5(1): 81 (1942)

Synonym: *Hygrophorus subaromaticus* (A.H. Sm. & Hesler) Largent, The Agaricales (Gilled Fungi) of California, 5. Hygrophoraceae (California): 106 (1985).

#### Holotypus

USA • California, Orick, Prairie Creek State Park, Nov 28 1937, (voucher <u>MICH: 10963</u> [Smith 9167]). Augmented here with ITS sequence of holotype: ZAP119. UNITE/PlutoF:<u>SH0910861.10FU</u> (1.5% threshold).

#### Description

**Pileus** 10–50 mm broad, convex to broadly convex with incurved margin, becoming plane or nearly so, sometimes with a low flattened umbo, colour “buffy brown” on the disc, “pale olive-buff” near the whitish margin (a dull olive grayish brown to pallid) in the type, pale Buff with or without a white margin in <u>iNaturalist:256214946</u> (ITS:<u>PV791511</u>), white with pale Buff disc in <u>iNaturalist:191417805</u> (ITS:<u>PP975575</u>) and pure white in both iNaturalist:9640562 (ITS:<u>MG926555</u>) and <u>iNaturalist:148999043</u> (ITS:<u>OR593569</u>), glabrous slimy-viscid, margin striatulate in the type but not in the recent collections. Context thin, very soft and fragile, whitish. **Stipe** 5-6 cm long, 6–10 mm thick concolorous with gills when fresh but drying pale grey like the pileus in type, equal, hollow, fragile, slimy viscid, as in *G. laetus*, glabrous. **Lamellae** bluntly adnate with decurrent tooth in type or sinuate, white, with a faint grey cast in type, broad sub-distant, edges even; 1–2 lengths lamellulae inserted. **Taste and odour** faint but disagreeable subaromatic in type, taste mild or slightly disagreeable, indistinct, similar to burnt plastic. Lamellae white (not fluorescing blue) under UVf365 nm. **Basidiospores** 8.1–10.2 × 5.6–6.1 µm in type description, pale yellow in Melzer’s reagent. Cuticle a thick (180–300 µm), gelatinous zone, the hyphae narrow colourless, more or less interwoven (an ixotrichodermium). **Hypodermium** a rather well-defined brownish zone. Pileus trama of radial hyphae. **Clamp connections** absent or very rare.

#### Habitat

On soil under coastal redwoods (*Sequoia sempervirens*).

#### Geographical distribution

Known only from coastal redwood forests in northwestern North America (Fig. 4B).

#### Additional specimens examined

USA • California: (1) Mendocino Co., Jackson State Forest, (39.3890,-123.68460, 26 Jan 2018, A. Rockefeller [voucher AR-2018a]), <u>iNaturalist:9640562</u> (ITS:<u>MG926555</u>); (2) Humboldt Co., Eureka Area, Arcata Community Forest, near *Sequoia sempervirens* (40.88326, -124.06329, 16 Feb 2023, Mandy Hackney [voucher HAY-F-000661]), <u>iNaturalist:148999043</u> (ITS:<u>OR593569</u>); (3) Humboldt Co., Eureka Area, McKay Community Forest, near *Sequoia sempervirens* (40.7501, -124.1446, 18 Nov 2023, Dean Lyons [voucher HAY-F-002303]), <u>iNaturalist:191417805</u> (ITS:<u>PP975575</u>).

#### Notes

This species, known only from the coastal regions of north California in North America, is found in coastal redwood (*Sequoia sempervirens*) forest. Its key distinguishing feature is its unpleasant odour but the odour is not always present. The lamellae appear white under UVf365 nm, distinguishing it from the bright blue fluorescence of lamellae in *G. calunus*. Though first recorded in 1937, most records date from 2013 or later. Reports of this species from Vancouver B.C. in Canada are a misapplied name. M. Berbee & F. Stonehouse at UBC obtained an ITS sequence of a collection by A. and U. Češka from Vancouver (UBC-F-3506) that most closely matched the type description of *H. subaromaticus*. The GenBank BLAST of that sequence confirmed that it had been misidentified as *H. subaromaticus* and instead belonged to a possibly unknown species, near *Cuphophyllus* sp. ‘PNW01’ and *C. subaustraligus*. Although an ixocutis was found on the pileus and stipe of the Vancouver collections, the specimens differed in having only a somewhat viscid rather than glutinous pileus that is typical of both *G. calunus* and *H. subaromaticus*.

### Key to *Gliophorus* sect. Unguinosae

|  |  |
| --- | --- |
| 1: Pileus pale, white with Buff tint or pale Drab | 2 |
| 1*: Pileus strongly pigmented, brown or greyish brown at least in centre | 4 |
| 2: Distributed in the UK, possibly Eurasian based on eDNA (pale forms of *G. irrigatus* rarely occurs, identifiable only by ITS sequence) | *G. alboviscidus* |
| 2*: Distributed in northwestern north America | 3 |
| 3: Lamellae blue under UVf365 nm, lacking any odour | *G. calunus* |
| 3*: Lamellae white under UVf365 nm, under coastal redwood and often with unpleasant burnt plastic or rubber odour | *G. subaromaticus* |
| 4: Distributed in Europe & the UK | *G. irrigatus* |
| 4*: Distributed in eastern North America | 5 |
| 5: Pileus centre dusky grey-brown or fuscous, lamellae broad | 6 |
| 5*: Pileus Drab Gray with darker striae to disc, faintly sulcate-striate to warm brown umbo; lamellae narrow, white; stipe long, slender (SH0910866.10FU; ITS: MZ269231; USA, NY) | iNat:30847408 |
| 6: Pileus margin straight when young, translucent-striate, rarely sulcate-striate, lamellae blue under UVf365 nm | *G. fumosus* |
| 6*: Pileus margin often inrolled & sulcate-striate; UVf365 nm unknown | *G. parafumosus* |

## Discussion

The first new species reported here, *G. alboviscidus*, is identical in morphology to *G. irrigatus*, except for its colour. Initial conclusions that it was an albino form were dispelled by ITS sequence analysis, which found that both specimens formed a distinct clade from European *G. irrigatus* s.s.. To our knowledge, Persoon did not provide any image of *G. irrigatus*, nor has any voucher survived, but it is likely that the specimen described in his monograph was found near Gottingen, Germany. Therefore, we neotypify *G. irrigatus*, based on voucher CFMR DEN-21 (KF291084), assessed by David Boertmann in Lodge et al. (2014), from Denmark, ca. 600 km north of Gottingen.

There was historically some debate as to whether *G. irrigatus* and *G. unguinosus* were distinct species (Arnolds 1990; Boertmann 2010). The latter species was named by Fries, who also provided a painting based on a Swedish specimen. However, there is no evidence from either vouchers or eDNA data that more than one grey or brown species within *Gliophorus* Sect. Unguinosae exists in Europe.

Our designated neotype of *G. irrigatus* (ITS:<u>KF291084</u>) and holotype of *G. alboviscidus* diverge by >9% in their ITS sequences. It is noteworthy that *G.irrigatus* specimens from across Europe form a distinct clade hitherto not reported beyond Europe, even when eDNA data are assessed (single exception, see above), whereas *G. alboviscidus* has a broader distribution, being detected in eDNA datasets from China.

*G. irrigatus*-like basidiomes are widely reported from North America. However, our sequence analysis found these species to be quite distinct (ca. 85% identity for ITS locus) from their morphologically very similar European relatives. Here we describe two sibling species found across eastern North America, *G. fumosus* and *G. parafumosus*, that form two genetically distinct clades but having only subtle differences in the morphology of the pileus margin. eDNA sequences of *G. fumosus* extend from eastern to western USA whereas *G. parafumosa* appears to be restricted to eastern North America.

A similar white to Buff coloured species from the *G. irrigatus* complex is known from the western USA (*G. subaromaticus*), often characterised by an unpleasant odour and recently found to have white lamellae under UVf365 nm. Sequence data from the holotype confirmed that this to belongs to *Gliophorus* Sect. Unguinosae. Also from the western US, a sixth species was found with pale or often more strongly pigmented basidiomes and bright blue lamellae under UVf365 nm, *G. calunus*. It is also clear from the phylogenetic analysis of sequence data that additional species exist within *Gliophorus* Sect. Unguinosae beyond the six species named here (e.g. OU003418 from Ecuador and MZ269231 from NY). In most cases, these are represented only by eDNA sequences, one exception being a single voucher (<u>iNaturalist:30847408/MZ269231</u>) from northeastern USA, which appears to be morphologically distinct and is included in our key.

The other exception is a southeastern US clade represented by a single voucher (<u>iNaturalist:251773868/PV223767</u>) provisionally named *Gliophorus* sp. ‘FL03’ from near Orlando, Florida that formed a clade with an environmental sequence from the Duke Forest in Durham, NC (GenBank:<u>AY969860</u>) and an ITS2 sequence from an ectomycorrhiza in Arkansas (GenBank:<u>MF665104</u>)(Fig. 2). This *Gliophorus* sp. ‘FL03’ appears inside a strongly supported clade with *G. alboviscidus* from Eurasia and *G. calunus* from northwestern North America, which suggests it is part of a lineage that was previously widespread and diverged into three Grayan disjunct species.

Lastly, the grey-brown species found in Australia and named *Hygrocybe irrigatus* (Young 2005) (pp 114/128) is likely to be divergent and to represent a new species. GlobalFungi search using ITS1 sequence of *G. fumosus* did detect eDNA (95% ID) hits from soil samples collected in New South Wales and Tasmania (Bissett et al. 2016), suggesting that a distinct species with darker pileus and most closely related to *G. fumosus* is indigenous to Australasia (Suppdata 2).

It is often accepted as a rule of thumb that species whose ITS sequences diverge by >3% represent different distinct species (Blaalid et al. 2013; Kauserud 2023), though there are examples of members of the same species having greater levels of ITS divergence, up to 6% (Hughes et al. 2013). In our study, the most closely related (named) species pair, *G. alboviscidus* and *G. calnus* diverge by ca. 4% but there is no evidence that their geographical ranges overlap even when eDNA data are assessed.

Assessing the distribution of novel species is by definition difficult. However, the advent and by now widespread use of eDNA metabarcoding provides a means to better assess species distributions. The GlobalFungi website provides an accessible means whereby such datasets may be interrogated and therefore provides an invaluable resource for assessment of species distributions; one which will become progressively more useful as eDNA metabarcoding datasets are published at an accelerating rate. One point important to note is that eDNA data cannot assess whether the sequences detected derive from mature individuals, an important element in fungal conservation (Ainsworth et al. 2013; IUCN 2024), where occurrence of basidiomes forms the basis of biological recording and conservation policy.

In the case of Hymenomycetes, which are mostly outcrossing, establishment of a primary mycelial (monokaryon) from single basidiospore must be followed by mating with a compatible basidiospore or primary mycelium to permit formation of the secondary mycelium (dikaryon) and later fruiting (James 2023). However, follow-on field surveys for fruitbodies of the species detected can be geo-focused based on eDNA data and this method has been successfully deployed in the UK (The Leasowes SSSI) to confer legal protection on sites of high grassland fungal diversity (Detheridge and Griffith 2021; UK_Government 2019).

Use of eDNA data to better understand the distributions of fungi has huge potential (Copo□ et al. 2024) but it does not (and cannot) define mature (reproducing) individuals (the basis of biological recording, conservation policy and e.g. GBIF). For agaric fungi, most form highly mobile uninucleate spores which may land and infrequently form mycelia at great distance. However, these ‘lost monokaryons’ will only proceed to form mature individuals if they mate with a compatible basidiospore or mycelium, a much rarer event than the establishment of the original mycelium, if far beyond the normal range of the species. There are additional alternative explanations for eDNA being detected outside the range of agaric species known from spore-producing basidiomes.

Besides long-range dispersal via haploid basidiospores forming monokaryotic mycelia that fail to encounter an opposite mating type, as noted above, we hypothesize three additional reasons why the distributions of eDNA sequences may exceed the known ranges of *Gliophorus* species based on basidome specimen records: 1) possible gaps or insufficient effort in collecting basidiomes or failing to collect basidiomes believed to be a common species (most likely explanation in our opinion); 2) spores may have dispersed beyond climatic zones that are currently or rarely conducive to fruiting; and 3) eDNA distributions exceeding the known range of basidiomes may indicate remnant populations of previously widespread distributions.

In the case of *G. irrigatus* s.s., basidiomes and eDNA have been intensively documented for Europe, including the UK and southern Scandinavia, regions where no other species are easily confused with it, basidiome and eDNA distributions are largely congruent (Figs. 3B & C). In contrast, the abundance of eDNA sequences of *G. alboviscidus* in eastern China, a species only known from basidiomes in the UK, might indicate insufficient surveys in eastern Eurasia, previous confusion of this new species with *G. irrigatus* from Europe, or a formerly widespread distribution that has become disjunct with changes in climate. The relatively small divergence in ITS sequences between Eurasian *G. alboviscidus* and northwestern North American *G. calunus* suggests disjunction of a formerly widespread range.

The new eastern North American species, *G. fumosus*, is abundantly documented by sequenced specimens and eDNA records in eastern North America as far west as Wisconsin while eDNA records extend west to the northern (Montana and Colorado) and southern (New Mexico) Rocky Mountains, plus the Cascade Mountains in western Oregon. The combined eDNA and basidiome distribution of *G. fumosus*, omitting the Cascades, is largely concordant with the distribution of basidiomes of *Chromosera lilacifolia* (Peck) (Grootmyers et al. 2025).

Taken together, this suggests that the widespread distributions of these two predominantly eastern North American species of Hygrophoraceae may represent remnants of a formerly more widespread distributions, but they differ in that *C. lilacifolia* is able to reproduce in the western part of its range whereas *G. fumosus* has either failed to reproduce or more likely only reproduces rarely beyond the midwestern USA and adjacent Canada and/or has been ignored. Occurrences in the mountains of New Mexico are concordant with a southern east-west dispersal pathway via the transvolcanic belt in northern Mexico that has been documented for several species and sister species of boletes (Ortiz-Santana et al. 2007). Grayan disjunction of sister species and genera between North American and east Asian taxa occurred 3-15 (-25) MYA (Petersen and Hughes 2007; Wang et al. 2004; Yih 2012), and the uplift of the Rocky Mountains caused changes in rainfall to their east resulting in extirpations, followed by Quaternary glaciation cycles that extirpated additional populations in parts of North America and western Europe (Yih 2012). The estimated divergence times for the Hygrophoraceae and *Gliophorus* are early enough to be consistent with this hypothesis. The Stem Age for the Hygrophoraceae was estimated as 125 MYA, and the divergence of the genus *Gliophorus* was dated to around 60-65 MYA (He et al. 2019).

Whilst there is evidence from eDNA that *G. subaromaticus* exists beyond North America, no fruitbodies have been observed beyond the far west of North America. However, the report of a subspecies of *G. irrigatus* (*Gliophorus irrigatus* f. *ammoniacus*) from Saarland, Germany, with “strong nitrous odour” (Schmitt 2022) suggests that *G. subaromaticus* may be more widely distributed. However, no sequence data nor any other details are published for this specimen. (<u>IF/MB#842679</u>), so potential synonymy cannot currently be resolved. eDNA studies at Aberystwyth University have detected *G. subaromaticus* in soil samples from upland (non-wooded) habitats in south Wales and the English Peak district (GWG unpub. data) but as noted above, these may well not represent mature individuals.

This study highlights how sustained efforts by citizen scientists—particularly through the Pembrokeshire Fungus Recording Network, with whom our first author is affiliated—can materially advance fungal systematics. Community-led observations and collections, integrated with molecular analyses, were essential to recognizing the novel diversity reported here; continued support for such networks will accelerate future taxonomic discovery. The Fungal Diversity Survey (Grootmyers et al. 2025; Ness 2021), and citizen scientists collecting and sequencing specimens through The MycoMap Network, Mycota Lab, the Ohio Mushroom DNA Lab, the Counter Culture Lab in Oakland CA, and sequencing of select foray specimens by the North American Mycological Association have been critically important in documenting new species, describing variation within species, and extending ranges of known species in North America.

In this study, we have been careful to obtain where possible holotype samples for ITS sequencing, and where this was not possible to neotypify species according to best practice guidelines. Failure to follow such an approach when naming new species can lead to future confusion and we advocate that fellow fungal taxonomists focus not only on naming new taxa but also clarifying the concepts of well-established species.

## Supporting information

Suppdata1

Suppdata2

Suppdata3

**Suppdata 1**. Details of all the vouchers and DNA sequences used in this study

**Suppdata 2**. The final alignment and trees (partitions, models, alignments and trees for both IQTREE and MrBayes runs)

**Suppdata 3** Phylogenetic reconstruction of the *Gliophorus* Sect. Unguinosae (ML tree) focused on ITS1 sequences, with *Gliophorus sciophanus* as outgroup. Numbers at salient nodes indicate % ultrafast bootstrap support (3000 replicates). The clade shown in red, adjacent to *G. fumosus*, is represented by two soil eDNA sequences from Australia (as detailed in Suppdata 1).

## Acknowledgements

The citizen science project in the UK was supported by The British Mycological Society’s Field Mycology DNA Barcoding programme. Citizen science in the USA for documenting and sequencing specimens was supported by the California Fungal Diversity Survey, Mycota Lab, The MycoMap Network, Ohio Mushroom DNA Lab, the North American Mycological Association Foray program in collaboration with the Field Museum of Natural History, and a US National Science Foundation grant for The Great Smoky Mountains National Park All-taxa Biological Inventory – Fungal Twig to K.W. Hughes at the University of Tennessee. Funding for eDNA sequencing in the USA was provided by National Science Foundation Grant DEB-2106130 Macrosystems Biology and NEON-Enabled Science Grant DEB-2106130 Macrosystems Biology and NEON-Enabled Science to B.A. Roy, A. Wilson, A.E. Arnold, J. Diez, M.E. Smith, P.G. Kennedy and S. Frey. We also thank PIs B.A. Roy, A. Wilson, M.E. Smith P.G. Kennedy and S. Frey for taking field samples of soil and litter, and H. Dawson for sequencing and C. Delevich for bioinformatics at University of Oregon. CALD is funded by the Brazilian National Council for Scientific and Technological Development (CNPq 446124/2024-9). We acknowledge the support of the Supercomputing Wales project, which was part-funded by the European Regional Development Fund (ERDF) via Welsh Government.

We are grateful to Caron Evans (IBERS Genomic Lab, Aberystwyth) for sequencing UK specimens and to Martyn Ainsworth, Andrew Detheridge, Matt Wainhouse, David Hawksworth and Paul Kirk for invaluable nomenclatural advice. We are grateful to Jorinde Nuytinck (Naturalis, Leiden), Åsa Kruys (Museum of Evolution (UPS), Uppsala University and Mats Wedin (Swedish Museum of Natural History, Stockholm) for confirming the absence of any relevant specimen or illustrations at their respective institutions. We thank M. Berbee, F. Stonehouse, and J. Ngo at the University of British Columbia in Canada for sequencing a putative *Hygrophorus subaromaticus* specimen at UBC collected by A. & O. Češka, and the following herbarium curators for assistance with loans: K. Golinsky at UBC, W. Gaswick & C. Christian at F. We thank B. Perry, Director of HAY, for arranging for W. Cardimona measure spores and observe clamp connections from two *G. calunus* collections.

