## Supplementary material for "Revisions to the *Gliophorus irrigatus* complex (Agaricales, Hygrophoraceae, *Gliophorus*, section Unguinosae) including a new waxcap, *G. alboviscidus*, from the UK but detected globally via soil eDNA, two new species from eastern North America, *G. fumosus* and *G. parafumosus*, plus a new species": Suppdata3

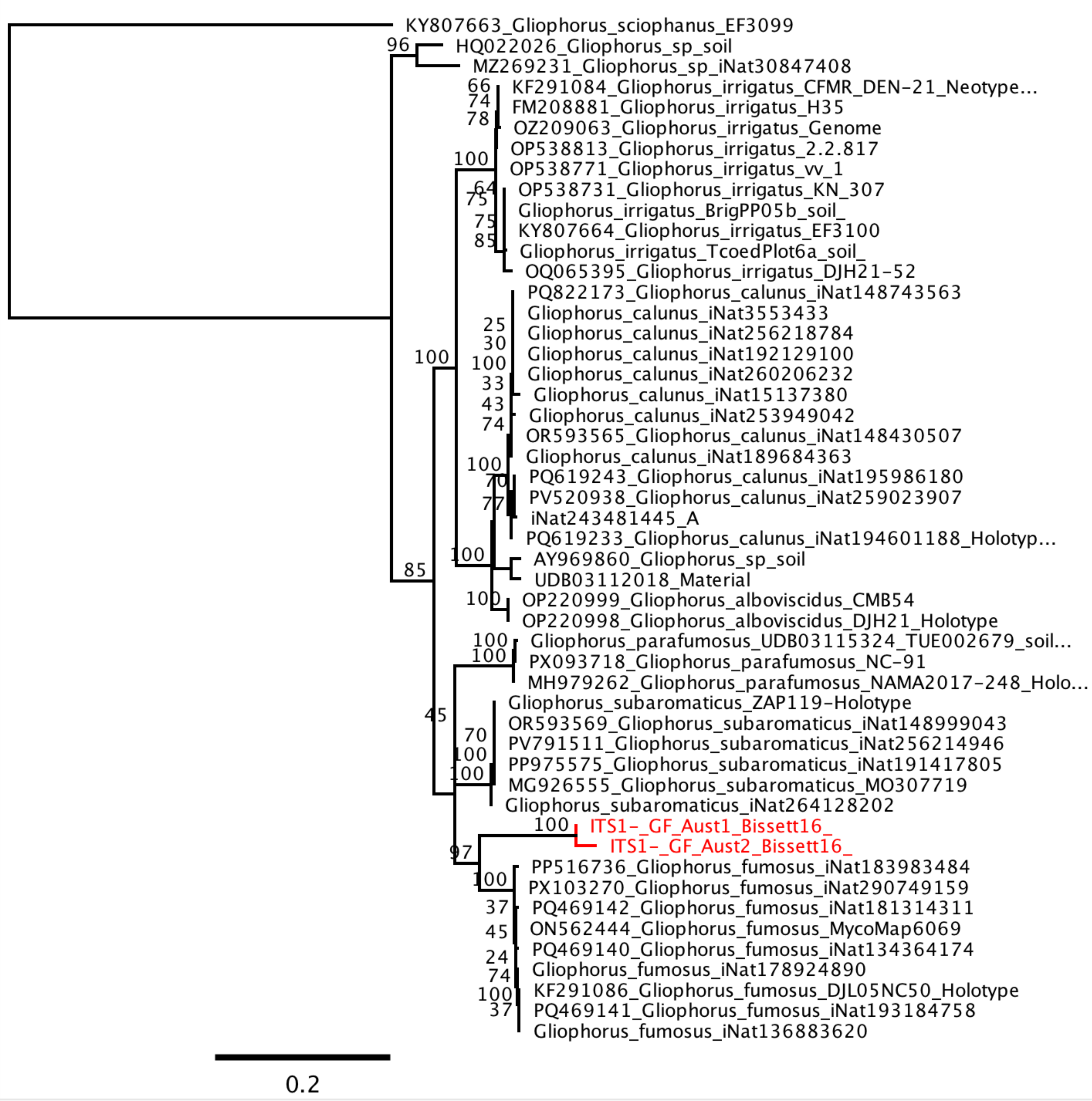


**Suppdata 3.** Phylogenetic reconstruction of the *Gliophorus* Sect. Unguinosae (ML tree) focused on ITS1 sequences, with *Gliophorus sciophanus* as outgroup. Numbers at salient nodes indicate % ultrafast bootstrap support (3000 replicates). The clade shown in red, adjacent to *G. fumosus*, is represented by two soil eDNA sequences from Australia (as detailed in Suppdata 1).
